# Visualising Cholesterol in Brain by On-Tissue Derivatisation and Quantitative Mass Spectrometry Imaging

**DOI:** 10.1101/2020.11.06.369447

**Authors:** Roberto Angelini, Eylan Yutuc, Mark F Wyatt, Jillian Newton, Fowzi Adam Yusuf, Lauren Griffiths, Benjamin Jordan Cooze, Dana El Assad, Gilles Frache, Wei Rao, Luke B. Allen, Zeljka Korade, Thu TA Nguyen, Rathnayake AC Rathnayake, Stephanie M Cologna, Owain W Howell, Malcolm R Clench, Yuqin Wang, William J Griffiths

**Author notes:** Corresponding author: William J Griffiths, **Email:**.

## Abstract

Despite being a critical molecule for neurobiology and brain health, mass spectrometry imaging (MSI) of cholesterol has been under reported compared to other lipids, due to the difficulty in ionising the sterol molecule. In the present work we have employed an on-tissue enzyme-assisted derivatisation strategy to improve detection of cholesterol in brain tissue sections. We report distribution and levels of cholesterol across specific brain structures of the mouse brain, in a model of Niemann-Pick type C1 (NPC1) disease, and during brain development. MSI revealed how cholesterol changes during development and that in the adult is highest in pons and medulla of the brain stem. Cholesterol was significantly reduced in the corpus callosum and other brain regions in the *Npc1* null mouse, confirming hypomyelination at the molecular level. Our study demonstrates the potential of MSI to the study of sterols in neuroscience.

## Introduction

Cholesterol is the most abundant individual molecular specie in plasma membranes of animals (Steck & Lange, 2018) accounting for approximately 20 - 25% of the lipid molecules in the plasma membrane of most cells (Dietschy & Turley, 2004), with only a small proportion of cellular cholesterol embedded in organelle bilayers (van Meer *et al*., 2008). Within membranes, cholesterol influences bilayer fluidity and permeability as well as lipid and protein sorting in membrane trafficking (Simons & Ikonen, 2000, Maxfield & van Meer, 2010). In brain, cholesterol makes up about 15% of the dry weight of white matter (WM) and is a major component of myelin sheaths that protect and insulate neuronal axons and are required for the fast transduction of neuronal signals (Brady *et al*., 2012, Lajtha *et al*., 2010). However, to date, little is known about how sterol concentrations vary in different anatomical locations or at sites of focal pathology (Almeida *et al*., 2015).

Cholesterol is metabolised to oxysterols, steroid hormones and bile acids (Russell, 2003). These metabolic pathways are at least partially operative in brain and their metabolic products and intermediates serve as regulatory and biologically active signalling molecules (Griffiths & Wang, 2019). In light of this, it is not surprising that impairment in sterol homeostasis and signalling have been implicated in a number of human disorders including neurodegenerative and neurodevelopmental conditions (Tabas, 2002, Björkhem *et al*., 2010, Porter & Herman, 2011). Dysregulation of cholesterol homeostasis has been implicated in Alzheimer’s disease (Larsson & Markus, 2018, Picard *et al*., 2018) and multiple sclerosis (Chataway *et al*., 2014), whilst inborn errors of cholesterol metabolism and transport can result in neurological disorders (Kanungo *et al*., 2013), such as Smith-Lemli-Opitz syndrome (SLOS, 7-dehydrocholesterol reductase deficiency) and Niemann-Pick disease types C1 and C2 (NPC1 and NPC2), respectively.

Traditionally, cholesterol analysis in tissue begins with homogenisation followed by lipid extraction, leading to loss of spatial information. To better understand sterol biochemical and physiological roles there is a need to match molecular abundance with exact location. To this end, careful dissection of specific brain regions can be coupled to classical gas chromatography (GC) – mass spectrometry (MS) and to liquid chromatography (LC) – MS (Quan *et al*., 2003, Heverin *et al*., 2004, Mast *et al*., 2017). An alternative method to map sterol concentrations in brain is by exploiting mass spectrometry imaging (MSI). For example, time-of-flight (ToF)-secondary ion MS (SIMS) is an MSI technique where cholesterol has been detected with high intensities, even at subcellular resolutions. However, a significant drawback with this approach is that ToF-SIMS is a surface sensitive technique and cholesterol has been shown to migrate to and crystallize at the surface, covering up all co-localizing species in the tissue. In the other hand, matrix-assisted laser desorption/ionisation (MALDI)-MSI has been employed to detect and identify multiple molecular species and simultaneously map their distribution in tissues sections (Caprioli *et al*., 1997, Norris & Caprioli, 2013, Spengler, 2015). It can generate pixelated MS data at near-cellular resolution providing spatial mapping of protein, peptide and lipid molecules according to X-Y position on a tissue section (Niehaus *et al*., 2019, Dreisewerd, 2014, Rompp & Spengler, 2013, Berry *et al*., 2011). MALDI-MSI has been used to image lipids in brain (Trim *et al*., 2008, Hankin *et al*., 2011, Woods & Jackson, 2006), however, cholesterol and other sterols tend to be poorly ionised by conventional MALDI and are discriminated against compared to other lipid classes that are more abundant. Cholesterol has been detected in MALDI-MSI studies (Tobias *et al*., 2018), but to enhance ionisation, other desorption methods have been employed, including nanostructure-initiator MS (Patti *et al*., 2010), sputtered silver-MALDI (Dufresne *et al*., 2013, Xu *et al*., 2015) and silver nanoparticle-MALDI (Roux *et al*., 2016, Muller *et al*., 2017). Silver ions coordinate with carbon-carbon double bonds providing cationic adducts of sterols in the MALDI matrix. Recently, “MALDI-2”-MSI has been developed, where a post-desorption second tuneable laser has been shown to enhance the ionisation of neutral lipid species including cholesterol, allowing improved visualisation in tissue sections (Barré *et al*., 2019, Soltwisch *et al*., 2015). Alternatively, derivatisation strategies can be utilised to enhance sterol ionisation. For in-solution studies we and others have exploited enzyme-assisted derivatisation for sterol analysis (EADSA) (Griffiths *et al*., 2013, Griffiths *et al*., 2016, Crick *et al*., 2015, Solheim *et al*., 2019) where the sterol molecule is reacted first with cholesterol oxidase enzyme in order to oxidise the 3β-hydroxyl group to a 3-oxo and then with Girard-P (GP) hydrazine to give a charge-tagged sterol hydrazone (Figure 1). This strategy enhances MS signal and provides unique structural information upon multistage fragmentation (MS^n^) which, together with retention time and accurate mass measurements, can provide unambiguous identification, even of isomeric species. Of note, others have exploited a similar but different derivatisation strategy to visualise by MSI endogenous and synthetic steroids, already possessing an oxo function, using Girard-T hydrazine (Cobice *et al*., 2016, Barré *et al*., 2016, Cobice *et al*., 2013).

**Figure 1.**
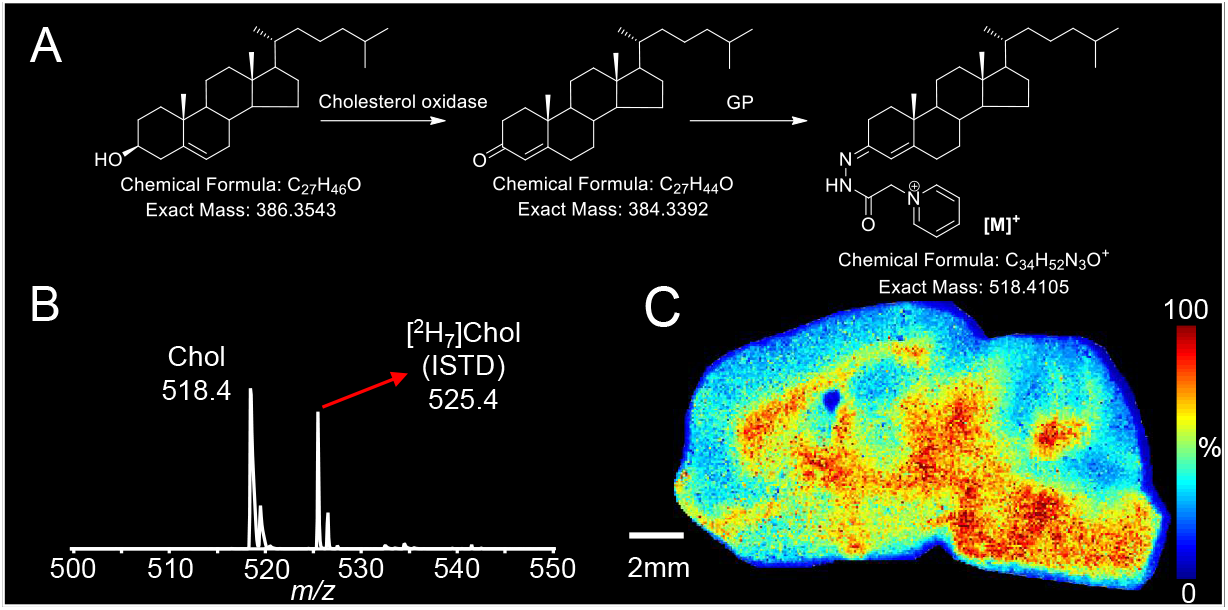
MSI of cholesterol in WT mouse brain via on-tissue enzyme-assisted derivatisation for sterol analysis (EADSA). (A) The on-tissue EADSA process using GP reagent occurs in two steps, both proceeding at 37°C and at ambient pressure in a humid atmosphere. The first step requires a water humidified atmosphere and is catalysed by the enzyme cholesterol oxidase, which converts the 3β-hydroxy-5-ene group to a 3-oxo-4-ene. Once the 3-oxo-4-ene group is in place reaction with GP hydrazine proceeds rapidly in a humid atmosphere above a solution of 50% methanol containing 5% acetic acid to give a GP hydrazone. (B) A typical mass spectrum generated in an EADSA-MALDI-MSI experiment for a single pixel. The spectrum, in the *m/z* range 500 - 550, is dominated by the signals from endogenous cholesterol and of sprayed-on standard [^2^H_7_]cholesterol. *In each pixel*, the peak at *m/z* 518.4 belonging to GP-derivatised brain cholesterol is normalised to the peak at *m/z* 525.5 belonging to GP-derivatised sprayed-on [^2^H_7_]cholesterol. By correlating the normalized intensity to a colour scale (%), an MS Image of the distribution of cholesterol across the mouse brain tissue section is created as shown in (C). (C) The MS Image is a heat map of a sagittal mouse brain tissue section showing the relative areal density of GP-derivatised cholesterol determined by MSI. MSI data were acquired on a vacuum-MALDI-TOF mass spectrometer. Data, normalized against sprayed-on [^2^H_7_]cholesterol, are shown using a “jet” scale. Scale bar 2 mm. Images were acquired at a pixel size of 50 µm. Isolation window width was 0.5 *m/z*.

We have adapted the EADSA protocol to MSI in order to image cholesterol in the developing and adult mouse brain as well as in a mouse model of NPC1 disease at 30 – 50 µm pixel size. We demonstrate the use of isotope-labelled standards to determine the absolute quantity of cholesterol in different anatomical regions of mouse brain. A quantitative MSI of the adult wild type (WT) mouse in sagittal sections was determined in the present study identifying pons and medulla of the brain stem as the regions with highest cholesterol level. The WT mouse was compared to the NPC1 mouse model which showed a significant reduction of cholesterol in the corpus callosum, caudate-putamen, thalamus, hypothalamus, midbrain, pons, and cerebellar WM. In the WT mouse brain at birth, we show that cholesterol is highest in the pontine hind brain that will develop into the cholesterol-rich pons region in the adult. The derivatisation-based method has the potential to be expanded to other low abundance sterols, while simultaneously detecting other lipid classes, visible without derivatisation. The EADSA-MSI technology shows potential in monitoring myelination processes and can be applied to the study of pathological diseases of the WM, including Alzheimer’s disease and multiple sclerosis, providing a new tool of investigation to neuroscientists.

## Results and Discussion

In MALDI-MS and electrospray ionisation (ESI)-MS, cholesterol is poorly ionised and is often detected as the ammonium adduct [M+NH_4_]^+^ at *m/z* 404.39 or as the dehydrated protonated molecule [M+H-H_2_O]^+^ at *m/z* 369.35 (McDonald *et al*., 2007). In lipidomics studies, these peaks can be distinguished from isobaric chemical noise by LC-MS. However, in MSI studies where orthogonal separations are not normally available, different strategies have been put in place to enhance the ionization efficiency of “intact” cholesterol in order to chemically resolve it from the background. Successful strategies include metal adduct formation, where Ag^+^ or Au^+^ coordinate with carbon-carbon double bonds providing cationic adducts of sterols in the MALDI matrix (Cologna, 2019). To further enhance ionization and desorption of sterols, here we exploit the EADSA method, previously used for in-solution analysis of sterols. Once the sterol analyte is specifically and effectively charge-tagged by EADSA (Figure 1A), it is readily analysed by MSI, thereby allowing its detection and identification (e.g. by MS^3^) and the mapping of its distribution. The advantage of this methodology is four-fold in that it (i) greatly increases sensitivity, (ii) allows for absolute quantification, (iii) enhances structural information, and equally importantly, (iv) increases analytical specificity. Here we report how EADSA has been adapted to work on brain tissue sections for MSI studies.

### Quantitative MSI of Sterols in WT Mouse Brain Reveals Regional Differences in Cholesterol Content

Initial studies were performed using sagittal mouse brain sections with a MALDI-time-of-flight (TOF) instrument (ultrafleXtreme, Bruker, Bremen, Germany). The GP-tagged cholesterol gives an intense [M]^+^ signal, as does sprayed-on [^2^H_7_]cholesterol, and dominates the resulting mass spectrum (Figure 1B). An MS Image of cholesterol distribution, normalized *in each pixel* to [^2^H_7_]cholesterol sprayed-on standard, is shown in Figure 1C in the form of a spatial heat map.

To confirm the identity of the signals assigned to cholesterol, we separately carried out an MS^3^ ([M]^+^**→**[M-Py]^+^**→**, where Py corresponds to the pyridine component of the GP-tag, see Supplemental Figure S1) analysis of the peaks at *m/z* 518.41 (cholesterol) and *m/z* 525.45 ([^2^H_7_]cholesterol) using atmospheric pressure (AP)-MALDI on an Orbitrap MS, see Figure 2. In Supplemental Figure S1, structures of the major fragment ions observed in Figure 2 are described. The fragment ion at *m/z* 163 (*b_3_-28) is formed by cleavage of the A/B-ring and is devoid of the CD-rings and the side-chain. It is present in MS^3^ spectra of both cholesterol (Figure 2A) and [^2^H_7_]cholesterol (Figure 2B) authentic standards and can thus be exploited in a multiple reaction monitoring (MRM)-like experiment to confirm the location of cholesterol and [^2^H_7_]cholesterol, sprayed directly on-tissue, in each pixel. Figure 2E shows that the MRM transition 518.4**→**439.4**→**163 is essentially absent off-tissue, while most notably enriched in the midbrain, pons, medulla and WM tracts of the cerebellum. Conversely, Figure 2F shows that the MRM transition 525.5**→**446.4**→**163 is saturated off-tissue, while being quite evenly distributed on-tissue. This transition does show some variation on tissue as a consequence of matrix effects. Note, for MS^3^ applications current software does not allow automated normalisation of cholesterol signals to the sprayed-on internal standard.

**Figure 2.**
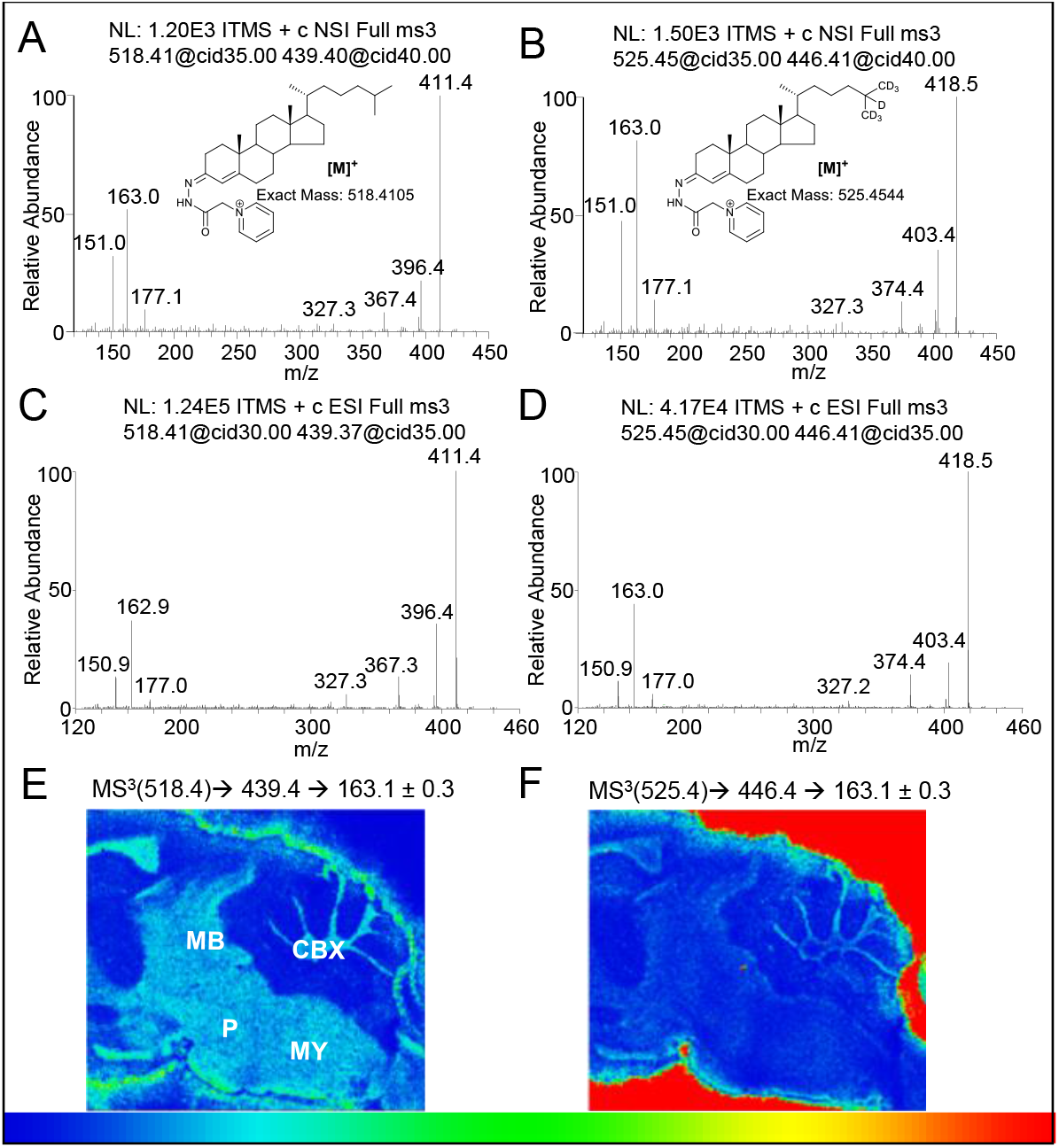
MS^3^ fragmentation patterns of (A) tissue-endogenous cholesterol and (B) sprayed-on [^2^H_7_]cholesterol, in a single pixel obtained in the LIT of an AP-MALDI-Orbitrap Elite instrument after EADSA on-tissue derivatisation of a brain tissue section and of (C) cholesterol and of (D) [^2^H_7_]cholesterol reference standards obtained in the LIT of an ESI-Orbitrap Elite instrument following in-solution EADSA. The MS^3^ spectra were obtained for the transitions [M]^+^**→**[M-Py]^+^**→**. (E, F) Images of the distribution of the MS^3^ fragment ion at *m/z* 163 from (E) cholesterol and (F) [^2^H_7_]cholesterol in a sagittal section of WT adult mouse brain.

Using MS^1^, areal densities were determined against a known density of sprayed-on internal standard in WT mouse brain sections, for selected brain structures (Figure 3A). The linearity of the on-tissue response of endogenous cholesterol versus the sprayed-on deuterated standard was determined by spraying eight consecutive tissue sections with [^2^H_7_]cholesterol at varying known densities (Supplemental Figure S2A). Examples of calibration curves obtained on whole-brain sections and considering the cerebellum as a region of interest (ROI) are shown in Supplemental Figure S2B & S2C, respectively. R^2^ for whole brain was determined to be 0.94 and for cerebellum 0.97. Our quantitative data reported in Figure 3B and in Table 1 (ng/mm^2^, mean over five biological replicates ± standard deviation) indicate that cholesterol abundance follows the pattern: pons (681.6 ± 123.9 ng/mm^2^) ≈ cerebellar white matter only (652.0 ± 119.8 ng/mm^2^) > medulla (613.8 ± 111.5 ng/mm^2^) > hypothalamus (545.7 ± 89.1 ng/mm^2^) ≈ mid brain (530.6 ± 70.6 ng/mm^2^) > corpus callosum (519.2 ± 55.9 ng/mm^2^) > thalamus (458.9 ± 59.2 ng/mm^2^) ≈ caudate-putamen (414.0 ± 74.1 ng/mm^2^) > whole cerebellum (395.0 ± 76.7 ng/mm^2^) > olfactory traits (348.5 ± 52.4 ng/mm^2^) ≈ cortex (327.8 ± 32.5 ng/mm^2^) ≈ hippocampus (326.3 ± 31.6 ng/mm^2^). Note that cholesterol was quantified via MSI in the whole cerebellum and in its WM tracts, whereas cholesterol content in cerebellar grey matter (GM) is estimated as described below. In previous reports (Quan *et al*., 2003), cholesterol synthesis and concentration were found to be higher in those regions of the central nervous system (CNS) containing heavily myelinated fibre tracts such as the brain stem (medulla, pons) and the midbrain. In contrast, concentration and turnover, were significantly lower in regions such as the cerebrum and whole cerebellum (Quan *et al*., 2003), which mainly represent GM structures with a reduced density of myelin. Our data in Figure 3 is consistent with these observations showing a significantly lower concentration of cholesterol in whole cerebellum and forebrain structures such as the cerebral cortex, olfactory tract, and hippocampus, as compared with midbrain or brain stem structures, such as medulla and pons. In an early report (Heverin *et al*., 2004), it has been shown that the concentration of cholesterol in pons is ∼2.5 times more than in cortex, which is also in agreement with our data showing 681.6 ± 123.9 ng/mm^2^ in pons and 327.8 ± 32.5 ng/mm^2^ in cortex (ratio 2.1). In earlier MSI studies, cholesterol was visualised in mouse brain in coronal or horizontal sections (Sjövall *et al*., 2004, Dufresne *et al*., 2013, Tobias *et al*., 2018). Sagittal MS Images have the advantage that the brain stem can be easily differentiated into the midbrain, pons and medulla regions (Figure 2E and Figure 3A). These brain stem structures show high cholesterol content (Figure 3, Table 1), in agreement with previous GC-MS and LC-MS studies (Quan *et al*., 2003, Heverin *et al*., 2004).

**Table 1:** Cholesterol areal density values in defined brain regions of WT and *Npc1*^*-/-*^ adult mice. In the present MALDI study n = 5 WT and n = 3 *Npc1*^*-/-*^ mice (10-weeks of age) were employed (three sections per mouse); in the LESA study n = 3 WT mice (12 weeks of age). Averages ± standard deviations are reported together with percentage agreement between MALDI- and LESA-MSI measurements of cholesterol levels in WT mouse brain, as are percentage differences in brain cholesterol levels as measured by MALDI MSI in WT and in *Npc1*^*-/-*^ mice.

| Brain area | MALDI MSI<br>WT<br>(ng/mm<sup>2</sup>) | LESA MSI<br>WT<br>(ng/mm<sup>2</sup>) | MALDI Vs LESA<br>WT<br>(% Agreement) | MALDI MSI<br>*Npc1*<sup>-/-</sup><br>(ng/mm<sup>2</sup>) | MALDI MSI<br>WT Vs *Npc1*<sup>-/-</sup><br>(% Difference) |
| --- | --- | --- | --- | --- | --- |
| olfactory traits | 348.5 ± 52.4 |  |  | 337.5 ± 35.2 | 3.2 |
| cortex | 327.8 ± 32.5 | 305.1 ± 53.8 | 93.1 | 307.4 ± 36.9 | 6.2 |
| corpus callosum | 519.2 ± 55.9 |  |  | 340.8 ± 44.6 | 34.4 |
| caudate-putamen | 414.0 ± 74.1 | 366.8 ± 4.7 | 88.6 | 357.5 ± 50.7 | 13.6 |
| thalamus | 458.9 ± 59.2 | 446.8 ± 51.7 | 97.4 | 352.6 ± 51.9 | 23.2 |
| hypothalamus | 545.7 ± 89.1 |  |  | 417.7 ± 50.2 | 23.5 |
| hippocampus | 326.3 ± 31.6 | 252.9 ± 27.3 | 77.5 | 288.9 ± 36.5 | 11.5 |
| mid brain | 530.6 ± 70.6 | 356.7 ± 74.8 | 67.2 | 383.1 ± 32.0 | 27.8 |
| pons | 681.6 ± 123.9 | 575.5 ± 122.0 | 84.4 | 454.2 ± 49.5 | 33.4 |
| medulla | 613.8 ± 111.5 | 421.3 ± 61.6 | 68.6 | 461.6 ± 111.3 | 24.8 |
| cerebellum | 395.0 ± 76.7 |  |  | 335.8 ± 40.0 | 15.0 |
| cerebellar white matter | 652.0 ± 119.8 | 509.2 ± 83.0 | 78.1 | 462.1 ± 58.7 | 29.1 |

**Figure 3.**
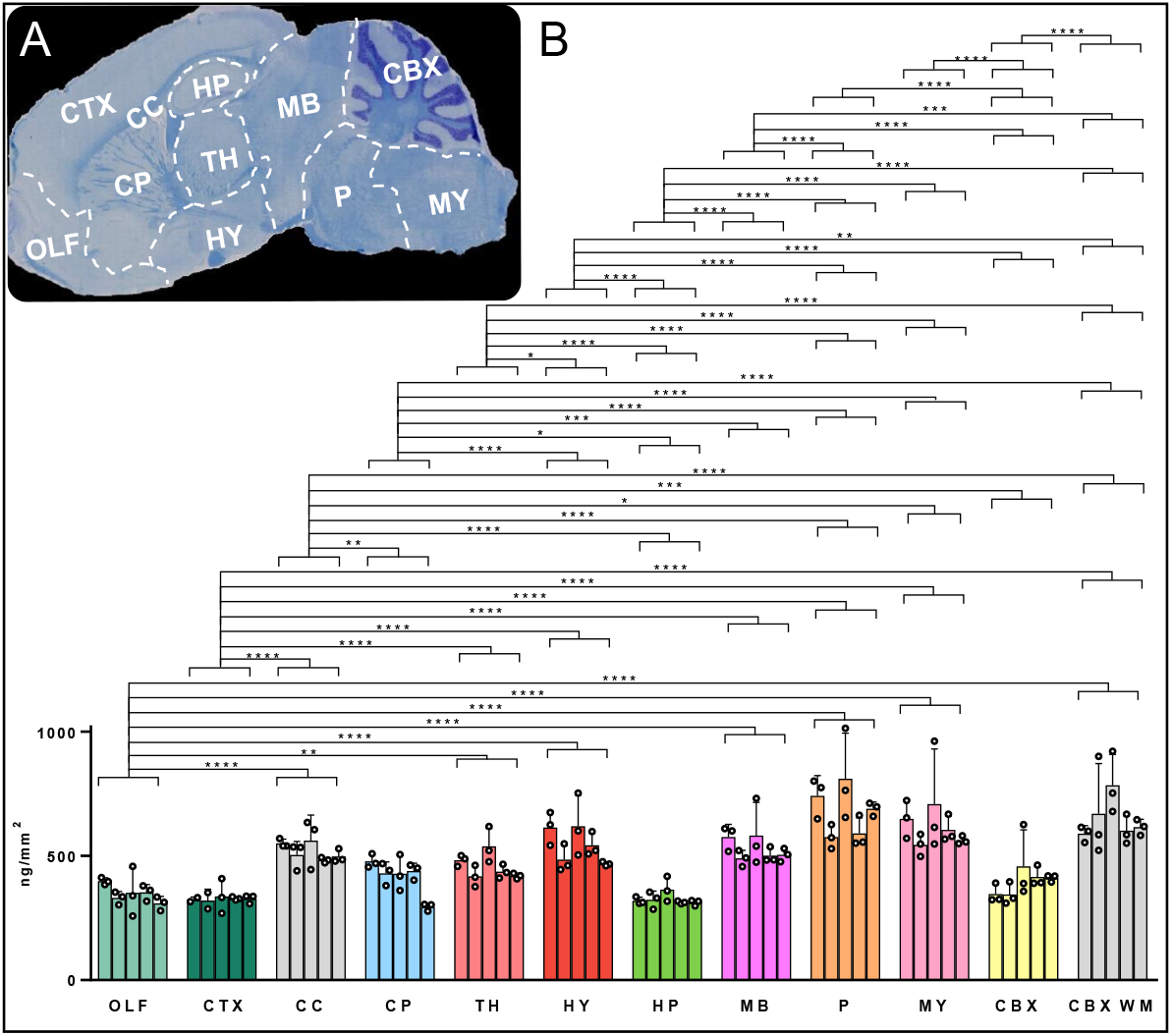
Quantitation of cholesterol in WT adult mouse brain via MSI. (A) LFB/CV staining for myelin of a sagittal mouse brain section adjacent to a section undergoing MSI. Major anatomical structures were identified by comparison with the corresponding Reference Atlas of the adult mouse brain provided by the Allen Institute of Brain Science (Lein *et al*., 2007) and are outlined with dashed lines: olfactory traits, OLF; cortex, CTX; corpus callosum, CC; caudate-putamen, CP; thalamus, TH; hypothalamus, HY; hippocampus, HP; midbrain, MB; pons, P; medulla, MY; cerebellum, CBX; cerebellar white matter, CBX WM. (B) Areal density (ng/mm^2^) of cholesterol in brain regions from five WT mice, averaged over different slices (see *Statistics* section). Significance levels from ANOVA using Tukey’s multiple comparison test are indicated for those brain regions showing significant differences. *P < 0.05, **P < 0.01, ***P < 0.001, ****P < 0.0001. Values for individual mice are given by separate histogram bars. The number of dots within each bar indicates the number of sections analysed for each mouse i.e. 3. Each dot within the bars correspond to region average for each brain slice. The height of each bar represents the mean of the region average for each mouse. The error bars indicate the SD of all the sections per mouse.

Interestingly, the distribution of gene transcripts of late-stage cholesterol biosynthetic enzymes match to regions of high cholesterol abundance i.e. midbrain, medulla, and pons regions. Please see mRNA expression data of [Dhcr24](https://mouse.brain-map.org/gene/show/50595), entrez ID 74754; [Dhcr7](https://mouse.brain-map.org/gene/show/13140), entrez ID 13360; and [Sc5d](https://mouse.brain-map.org/experiment/show?id=71724560), entrez ID 235293 provided by the Allen Mouse Brain Atlas (Lein *et al*., 2007). Of note, the abundance of cholesterol in the corpus callosum and in the fibre tracts of the caudate-putamen mirrors the distribution of transcripts unique to myelinating oligodendrocytes that wrap around axons during development. See [Mbp](https://mouse.brain-map.org/experiment/show?id=79632288), entrez ID 17196; [Plp1](https://mouse.brain-map.org/experiment/show?id=75496146), entrez ID 18823; [Cnp](http://developingmouse.brain-map.org/experiment/show/1176), entrez ID 12799 provided by the Allen Mouse Brain Atlas (Lein *et al*., 2007).

When MS^1^ data at high mass resolution is visualised in a peripheral sagittal section taken at a plane about 3 mm from the midline, the distribution of GP-derivatised cholesterol at *m/z* 518.4103 is clearly enhanced in specific regions of brain (Figure 4A). These are either WM tracts such as the corpus callosum and cerebellum, or brain regions (deep GM structures) containing myelinated fibres, such as the pons, medulla of the brain stem and caudate-putamen of the diencephalon. In Figure 4 the selected sagittal plane does not include midbrain but shows interesting hippocampal features described below.

**Figure 4.**
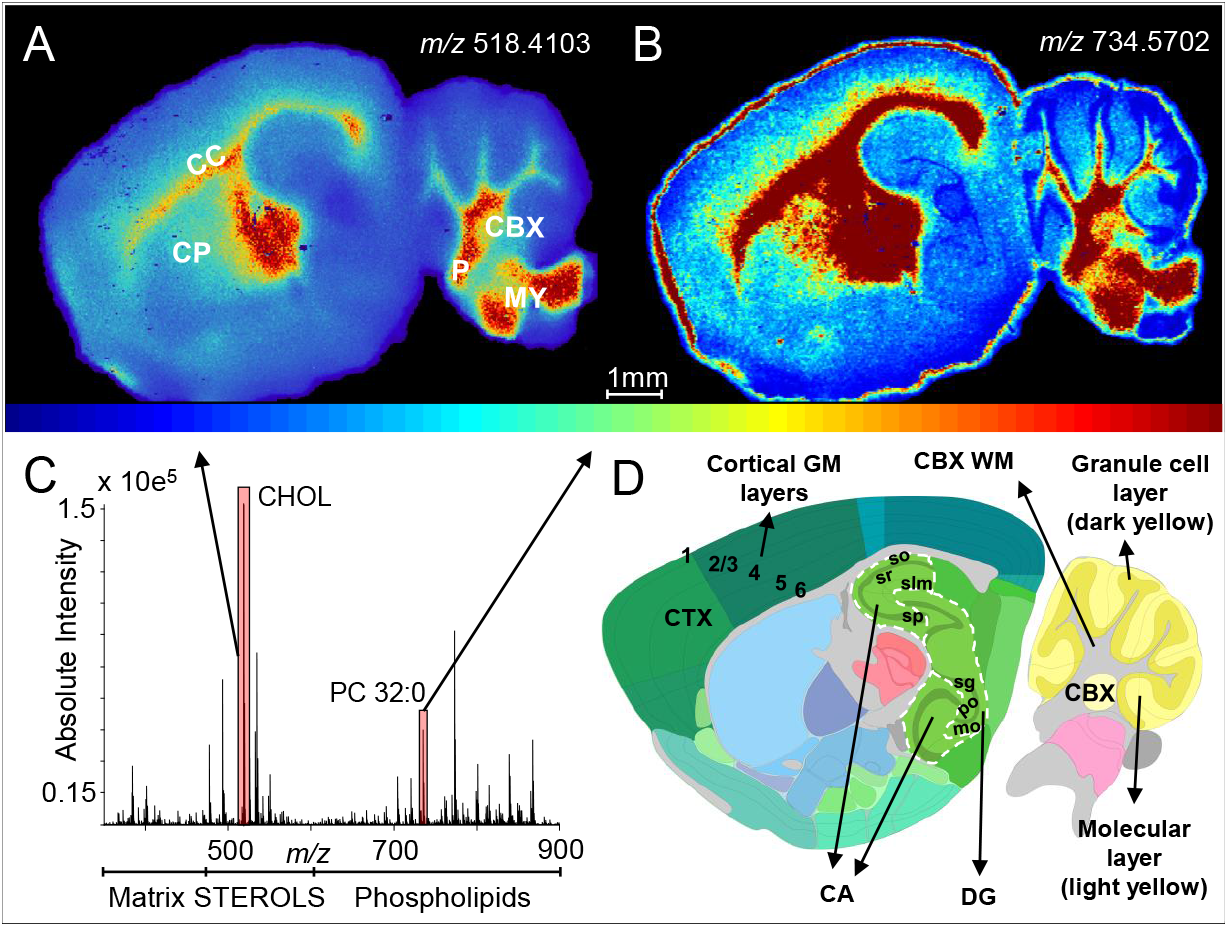
AP-MALDI-MSI at high mass resolution of cholesterol in sagittal sections of WT adult mouse brain. The data was obtained on Orbitrap instrument. (A) Distributional heat map of cholesterol. (B) Distributional heat map of a major structural phospholipid assigned to PC 32:0. (C) Typical AP-MALDI MSI spectrum averaged over the entire MSI dataset after on-tissue EADSA derivatisation showing sterol and phospholipid (and other brain lipids) signals that can be detected simultaneously. The spectrum contains hundreds of signals that can be mapped. (D) Anatomical layering of cortex, cerebellum and hippocampus. Layers of the cortex (CTX, dark green) and of the cerebellum (CBX, yellow) are shown. Cerebellar WM is in the lobules (coloured grey) of the cerebellum (CBX) which are surrounded by the granule cell layer (dark yellow) and then by the molecular layer (light yellow). Layers of the dorsal and ventral hippocampal formation are also visible in the selected sagittal plane in both Ammon’s horn (CA) and Dentate Gyrus (DG). The CA is layered into (dorsal to ventral) strata oriens (so), radiatum (sr) and lacunosum-moleculare (slm) (all in light green) with the pyramidal layer (sp, dark green) stratified between the strata oriens (so) and radiatum (sr). The DG shows the central granule cell layer (sg, dark green) in between the molecular (mo) and polymorph (po) layers (both in light green). Image credit Allen Institute for Brain Science: Adult Mouse, P56, Sagittal, Image 7 of 21 id=100883846 (Lein *et al*., 2007). In (A, B) isolation window width was 7 mmu, pixel size was 30 µm, mass deviation for cholesterol was <1 ppm using [^2^H_7_]cholesterol as a lock mass. Images normalized against sprayed-on [^2^H_7_]cholesterol.

Notably, using our method for on-tissue cholesterol derivatisation, and in contrast to ToF-SIMS, other lipids can be mapped simultaneously particularly when experiments are carried out under atmospheric pressure ionisation (i.e. AP-MALDI - Orbitrap and desorption-ESI (DESI) – Q-TOF). To show the potential of our approach, we report the MS image of PC 32:0 at *m/z* 734.5702 (about 1 ppm deviation from the theoretical *m/z*), normalized to [^2^H_7_]cholesterol sprayed-on standard (Figure 4B), but many other peaks could be similarly imaged. Please note that the peak at *m/z* 734.5702 could be similarly assigned to PE 35:0. However, phospholipids containing fatty acids with an odd number of carbon atoms are minor species in animals.

Interestingly in Figure 4A, a continuous gradient of cholesterol concentration is observed in the forebrain, going out from the corpus callosum, where cholesterol is at an areal density of about 520 ng/mm^2^, decreasing on moving through the overlying layers of the neocortex correlating with the higher myelination index of the corpus callosum, WM, in comparison to the cortical GM layers. Indeed, in the cerebral cortex, cholesterol was quantified to be about 330 ng/mm^2^ (Figure 4D shows reference anatomy). The continuous cholesterol gradient is mirrored by the distribution of the presumed phosphatidylcholine (PC 32:0) in the same brain regions (Figure 4B). Within the cerebellum, a structure with a central component of WM surrounded by partly myelinated GM, a decreasing concentration of cholesterol going from the WM of the cerebellum (CBX WM, 652.0 ± 119.8 ng/mm^2^) to the granule cell layer and molecular layer of the GM can be observed (Figure 4D shows reference anatomy). Note that cholesterol density in cerebellar GM can be estimated (CBX – CBX WM = CBX GM) to be about 260 ng/mm^2^. Here, the cholesterol smooth gradient contrasts with the step gradient shown by PC 32:0 (Figure 4B) which is deficient in the granule cell layer of the cerebellum but more evident in the molecular layer. Differential gradients for both cholesterol and PC 32:0 can be observed also in the Ammon’s horn and the dentate gyrus of the hippocampal formation. Figure 4B shows that PC 32:0 is deficient in the pyramidal layer of the Ammon’s horn and in the granule cell layer of the dentate gyrus while it is more evident in the strata oriens, radiatum and lacunosum-moleculare of the Ammon’s Horn as well as in the molecular and polymorph layers of the dentate gyrus (Figure 4D shows reference anatomy). The step gradient of PC 32:0 demonstrates the outlines of the Ammon’s horn and of the dentate gyrus, while the cholesterol distribution gradient is more continuous across these structures (Figure 4A). Indeed cholesterol, although being more concentrated in the myelin sheaths of the WM, is a key component of all membranes and, therefore, is present throughout the brain. Moreover, cholesterol is by far the major representative of its class, with other sterols present at comparatively low abundance and not exerting any known structural function in membranes. On the contrary membrane phospholipids such as PC, have high structural diversity (Harayama & Riezman, 2018) and may show both a continuous gradient (as in the cortex), or a step gradient (as in cerebellum and hippocampus) for an individual molecular specie, even within layers of the same brain structure.

The EADSA-MALDI-MSI quantitative assessment of cholesterol areal density in defined brain regions of the WT mouse can be compared with measurements previously obtained with a similar but different experimental approach exploiting low-spatial resolution (400 µm pixel size) Liquid Extraction for Surface Analysis (LESA) i.e. EADSA-LESA-LC-MS (Yutuc *et al*., 2020). Table 1 reports the values obtained and their standard deviations, in each defined brain region of WT mouse, for both the present and the EADSA-LESA-LC-MS study. The agreement between the studies is also reported. The agreement was > 90% for very homogenous regions such as cortex and thalamus, it was about 80% for caudate-putamen, hippocampus, pons and cerebellar white matter, and was > 67% for heterogeneous structures such as midbrain and medulla.

### Neuroimaging of Cholesterol in the Developing Mouse

To illustrate how our EADSA-MSI method can be used to monitor brain cholesterol distribution during development we compared tissues from mice at 1-day and 10-weeks. At birth, myelination is in its very early stage, while at 10 weeks is nearly completed (Goffinet & Rakic, 2000). During development, the cholesterol content in the whole brain goes from about 4 mg/g at birth up to about 15 mg/g in the adult at 26-weeks (Dietschy, 2009), and comes from local synthesis only (Dietschy & Turley, 2004). During the first three weeks of life, when myelin sheaths are being generated, the rates of cholesterol synthesis and accumulation in brain are high at about 250 µg/day (Quan *et al*., 2003), and drop rapidly beyond three weeks of age (Dietschy, 2009). Body growth is completed at about 7 weeks, with most strains reaching sexual maturity between 6 - 8 weeks. Therefore, we can use the arbitrary but reasonable date of postnatal week 10 as the definition of young adult mouse (Hedrich & Bullock, 2012).

EADSA-MSI was employed to visualise cholesterol distribution in the mouse brain at 1-day and at 10-weeks (Figure 5). We compared MSI of cholesterol with Luxol Fast Blue (LFB) chemical stain and Cresyl Violet (CV; as counterstaining) histology, a traditional but nonspecific histological stain for myelin (Kluver & Barrera, 1953), as illustrated in Figure 5C (newborn) and Figure 5D (10-weeks). Figure 5E shows the MSI of cholesterol distribution at 1-day around the time when oligodendrocytes start to contribute to cholesterol synthesis (Saher & Stumpf, 2015) and Figure 5F shows the distribution at 10-weeks.

**Figure 5.**
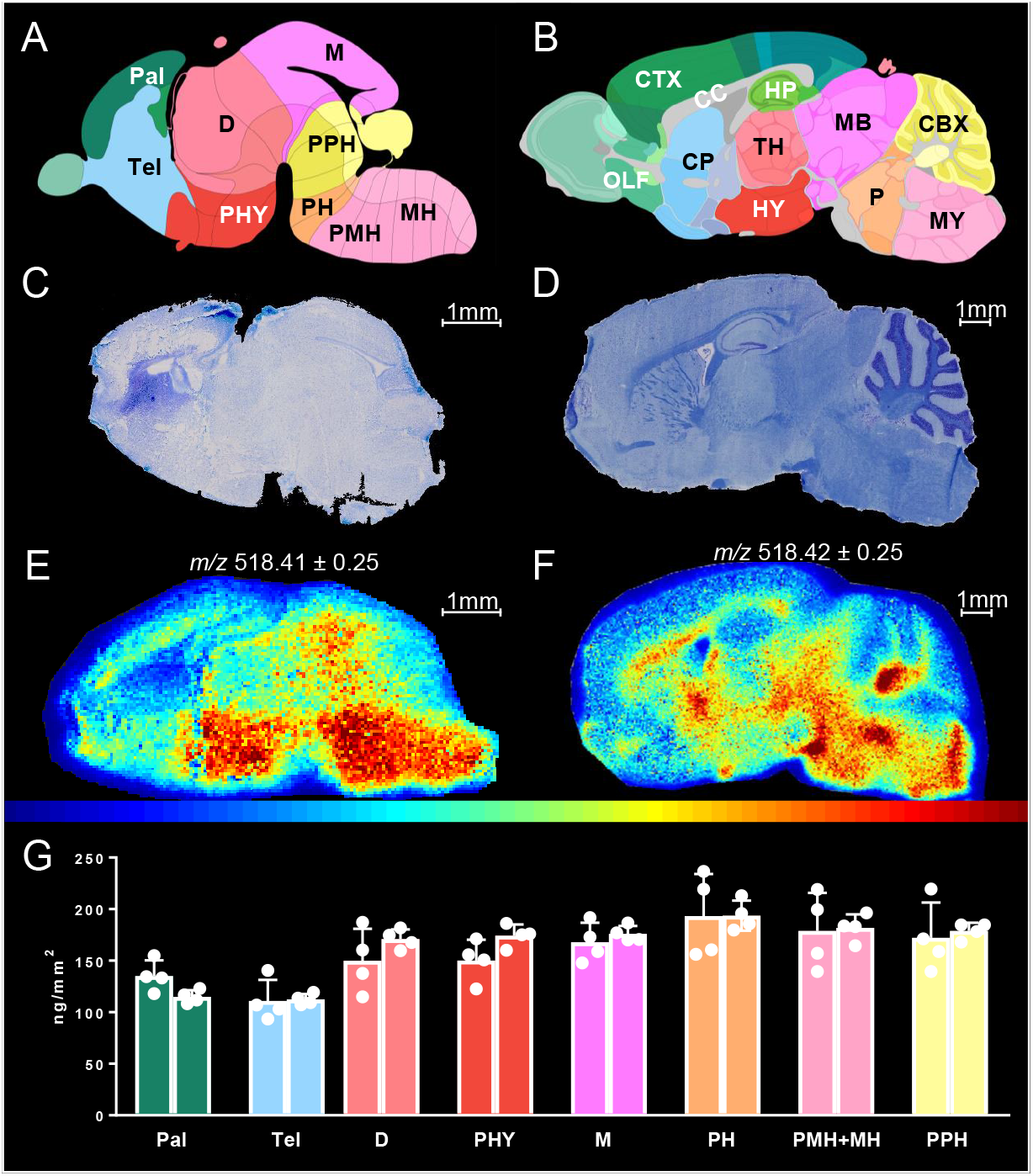
Quantitative MSI of cholesterol in the mouse brain at birth and at 10-weeks. In (A, C, E) 1-day-old newborn and in (B, D, F) 10-week-old adult mouse. (A, B) Images from the Allen Institute of Brain Science depicting mouse brain sagittal sections with annotations of anatomical structures. The 1-day-old newborn is matched with the E18.5-day embryo atlas image (A). Pallium, Pal; Telencephalic vesicle, Tel; Diencephalon, D; Peduncular Hypothalamus, PHY; Midbrain, M; Pontine Hindbrain, PH; Pontomedullary and Medullary Hindbrain, PMH+MH; Prepontine Hindbrain, PPH. (B) Adult mouse, abbreviations as in Figure 3. (C, D) LFB/CV staining of sagittal mouse brain sections adjacent to sections undergoing MSI. The newborn mouse brain appears pale indicating lack of myelination. (E, F) MSI of cholesterol after on-tissue EADSA by a vacuum-MALDI-MS. Data, normalized against sprayed-on [^2^H_7_]cholesterol, are shown using a “jet” scale. Scale bars 1 mm. Images were acquired at a pixel size of 50 µm. (G) Areal density (ng/mm^2^) of cholesterol in brain regions from two newborn WT mice, each averaged over 4 slices (see *Statistics* section). Values for individual mice are given by separate histogram bars. The number of dots within each bar indicates the number of sections analysed for each mouse i.e. 4. Each dot within the bars correspond to region average for each brain slice. The height of each bar represents the mean of the region average for each mouse. The error bars indicate the SD of all the sections per mouse. Image credit Allen Institute: Developing Mouse, E18.5, Image 16 of 19 id=100740373 and Adult Mouse, P56, Sagittal, Image 15 of 21 id= 100883867 (Lein *et al*., 2007).

As measured by quantitative EADSA-MSI, the newborn (Figure 5G, Supplemental Table S1) shows the highest cholesterol level in the pontine hindbrain (193.4 ± 28.4 ng/mm^2^) and in the medullary and pontomedullary hindbrain (180.3 ± 25.6 ng/mm^2^) that will develop into the cholesterol-rich pons and medulla of the adult mouse (Figure 3B) (Goffinet & Rakic, 2000). The lowest levels of cholesterol are detected in the telencephalic vesicle (Tel, 111.7 ± 13.9 ng/mm^2^) and in the Pallium (125.1 ± 15.3 ng/mm^2^) which will develop into the cortex, olfactory tracts, hippocampus and caudate-putamen (Goffinet & Rakic, 2000). Similar to the newborn, these are regions with low cholesterol in the adult (Figure 3, Table 1) except for the caudate-putamen which contains some fibre tracts in the adult that are not yet formed in the newborn (Goffinet & Rakic, 2000). The pro-hypothalamic region (peduncular hypothalamus), which begets the adult hypothalamus and associated fibre tracts, shows a diffused enrichment in cholesterol in the newborn (Figure 5A & E), while the hypothalamus in the adult accumulates cholesterol only in surrounding fibres (Figure 5B & F). A striking difference between 1-day and 10-week animals is the lack of a visible corpus callosum (CC) in the newborn. In the mouse, myelination of the CC is reported to begin at 11 days after birth (Sturrock, 1980) and CC is detected by histological methods at around 16-17 days of age (Wahlsten, 1984). In contrast to the newborn, in the 10-week adult the CC is fully formed (Hedrich & Bullock, 2012). In particular, in the adult the thicker regions of the CC show enrichment in cholesterol, namely the rostrum-genu (frontal, 504.3 ± 57.7 ng/mm^2^, see Supplemental Figure S3), the body (central, 546.2 ± 66.3 ng/mm^2^) and the splenium (back, 500.4 ± 35.3 ng/mm^2^), while the thinnest part of the CC, the isthmus connecting the body and the splenium is the CC structure with lowest cholesterol abundance (457.8 ± 51.2 ng/mm^2^). It is reassuring that the anatomical thickness of the CC matches with cholesterol enrichment as determined by EADSA-MSI.

Worthy of note, the unspecific LFB myelin stain of the newborn provides little distributional information as compared to the MSI heat map for cholesterol (Figure 5C cf. 5E). Indeed, by MSI a differential distribution of cholesterol can be detected across different areas of the newborn mouse brain, which does not match the distribution of myelin as assessed by LFB, which shows a very pale and uniform staining pattern across many regions in the newborn. Conversely the comparison of MSI and LFB in the adult shows an overlap of stain and mass spectral cholesterol signal intensity. Importantly, MSI is highly sensitive and as applied here, specific for cholesterol, while the exact molecular species bound by LFB remain uncertain (Blackwell *et al*., 2009). The distribution of cholesterol via MSI shows some overlap with gene expression data available at the Allen Brain Atlas (Lein *et al*., 2007) for lineage-specific markers of myelin-producing oligodendrocytes at the perinatal stage (E18.5). See for example expression of [Mbp](http://developingmouse.brain-map.org/experiment/show/100054687), entrez ID 17196; [Plp1](http://developingmouse.brain-map.org/experiment/show/100085380), entrez ID 18823; and [Cnp](http://developingmouse.brain-map.org/experiment/show/100077381), entrez ID 12799 in the mouse at E18.5. However, the first group of neurons in developing brain are monoaminergic neurons, see for example expression of [Th](http://developingmouse.brain-map.org/experiment/show/100071454), entrez ID 21823; [Tph2](http://developingmouse.brain-map.org/experiment/show/100057249), entrez ID 216343; and [Dbh](http://developingmouse.brain-map.org/experiment/show/100071430), entrez ID 13166 in the mouse at E18.5, and at P0 neurons provide their own source of cholesterol and it is likely that cholesterol synthesising and developing neurons contribute to the MS image of cholesterol in the newborn. Notably, the cholesterol distribution in the newborn mouse imaged by vacuum MALDI-TOF (Figure 5E) is consistent with the image of an adjacent brain section produced by DESI-Q-TOF (Supplemental Figure S4E), proving the robustness of the EADSA-MSI approach.

Finally, as measured by EADSA-MSI the whole-brain areal density of cholesterol in the newborn is about 160 ng/mm^2^ while it is about 480 ng/mm^2^ in the adult at 10 weeks, showing a 3-fold increase. Our data is in good agreement with previous reports (Quan *et al*., 2003) where the cholesterol content in the newborn was determined to be about 4 mg/g at birth and to increase to about 10 mg/g at 10 weeks, showing a 2.5-fold increase. In summary the present data demonstrate that EADSA-MSI can be used effectively to monitor cholesterol abundance in brain structures during development.

### Neuroimaging of Cholesterol in the Niemann-Pick Disease type C1 Shows a Lack of Cholesterol in Hypomyelinated Fibres Tracts

Niemann-Pick Disease, type C is a neurodegenerative, lysosomal storage disorder, characterized by accumulation of unesterified cholesterol and sphingolipids in the endo-lysosomal system (Vanier, 2015). The disease is caused by mutations in the encoding region of genes either for the lysosomal transmembrane protein, NPC1 (95% of cases) or the small cholesterol-binding soluble glycoprotein, NPC2 (∼4% of cases) (Loftus *et al*., 1997, Naureckiene *et al*., 2000). These two proteins work together to transport cholesterol through the late endosomal-lysosomal membrane into the cytosolic and metabolically active cholesterol pool. Patients with NPC disease show extensive hypomyelination that manifest in cerebral and cerebellar atrophy as well as WM hypoplasia as detected in MRI scans (Palmeri *et al*., 1994).

In the present study we analysed the cholesterol content and distribution in the brain of the *Npc1*^-/-^ mouse where the gene was knocked out by a naturally occurring retroposon-driven frameshift mutation (BALB/cNctr-*Npc1*^*m1N*^/J, https://www.jax.org/strain/003092) (Morris *et al*., 1982, Shio *et al*., 1982, Bhuvaneswaran *et al*., 1982). In the brain of this mouse, at the 7-week time point, cellular dysfunction translates into loss of many large neurons. In particular, Purkinje cells of the cerebellum are particularly sensitive to NPC pathology and are largely lost in patients (Gilbert *et al*., 1981) and in the mouse model (Higashi *et al*., 1993). Moreover, the brain of the *Npc1*^*-/-*^ mouse generally shows severe dysmyelination of fibre tracts with impairment of oligodendrocyte maturation (Saher & Stumpf, 2015). As in patients (Palmeri *et al*., 1994), oligodendrocyte loss and dysmyelination may result in hypoplasia of the corpus callosum in this mouse model (German *et al*., 2001, German *et al*., 2002). When NPC1/2 proteins are lacking, cholesterol and other lipids remain in the late endosomes/lysosomes and are not transported into the endoplasmic reticulum (ER) and, therefore, sterol homeostasis is undermined by the lack of feedback regulatory mechanisms, i.e. free cholesterol accumulates in the late endosomes/lysosomes compartment while the rest of the cell perceives a shortage of sterol. Indeed, in the *Npc1*^-/-^ mouse, sterol synthesis is not blocked by accumulating cholesterol as cholesterol overload is confined to lysosomes, consequently SREBP2 processing is constitutively active and target gene expression is maintained driving cholesterol synthesis (Liu *et al*., 2009). In this complicated landscape, it is important to understand the downstream effects of lysosomal cholesterol accumulation on defined brain areas, as well as on specific neuronal cell populations, *in vivo*. Previous work on lipid accumulation in NPC1 disease showed how several tissue are affected by changes in lipid composition (Zhou *et al*., 2011, Kulinski & Vance, 2007, Liu *et al*., 2000, Fan *et al*., 2013, Liu *et al*., 2009, Reid *et al*., 2008). However, in these studies tissue lysate lipidomics was employed to obtain sensitive, specific and quantitative data at the expense of information regarding spatial distribution. Recently one MSI study has assessed lipid changes in this *Npc1*^*-/-*^ mouse but was limited to the cerebellum (Tobias *et al*., 2018).

In the present study we have exploited MSI to study the whole brain. The chosen time point was of 10-weeks when the phenotype is severe but not yet lethal: *Npc1*^*-/-*^ mice die at an average of 84 days (12-weeks) (Liu *et al*., 2009). Figure 6A and 6B show the MSI spatial heat maps of cholesterol distribution in the WT and *Npc1*^-/-^ mouse brain, respectively. These heat maps can be compared with Figure 6C and 6D showing LFB/CV histological staining for myelin of corresponding adjacent brain tissue sections. For further comparison of MALDI-MSI with histology, the density of myelinated fibres in the caudate-putamen (Figures 6E and Supplemental S5A) and the number of Purkinje cells in the cerebellum (Figures 6F and Supplemental S5B) of WT and *Npc1*^-/-^ mouse were determined from the histological data obtained via LFB/CV staining. Figure 6G shows cholesterol levels in selected brain ROI as quantified by MSI.

**Figure 6.**
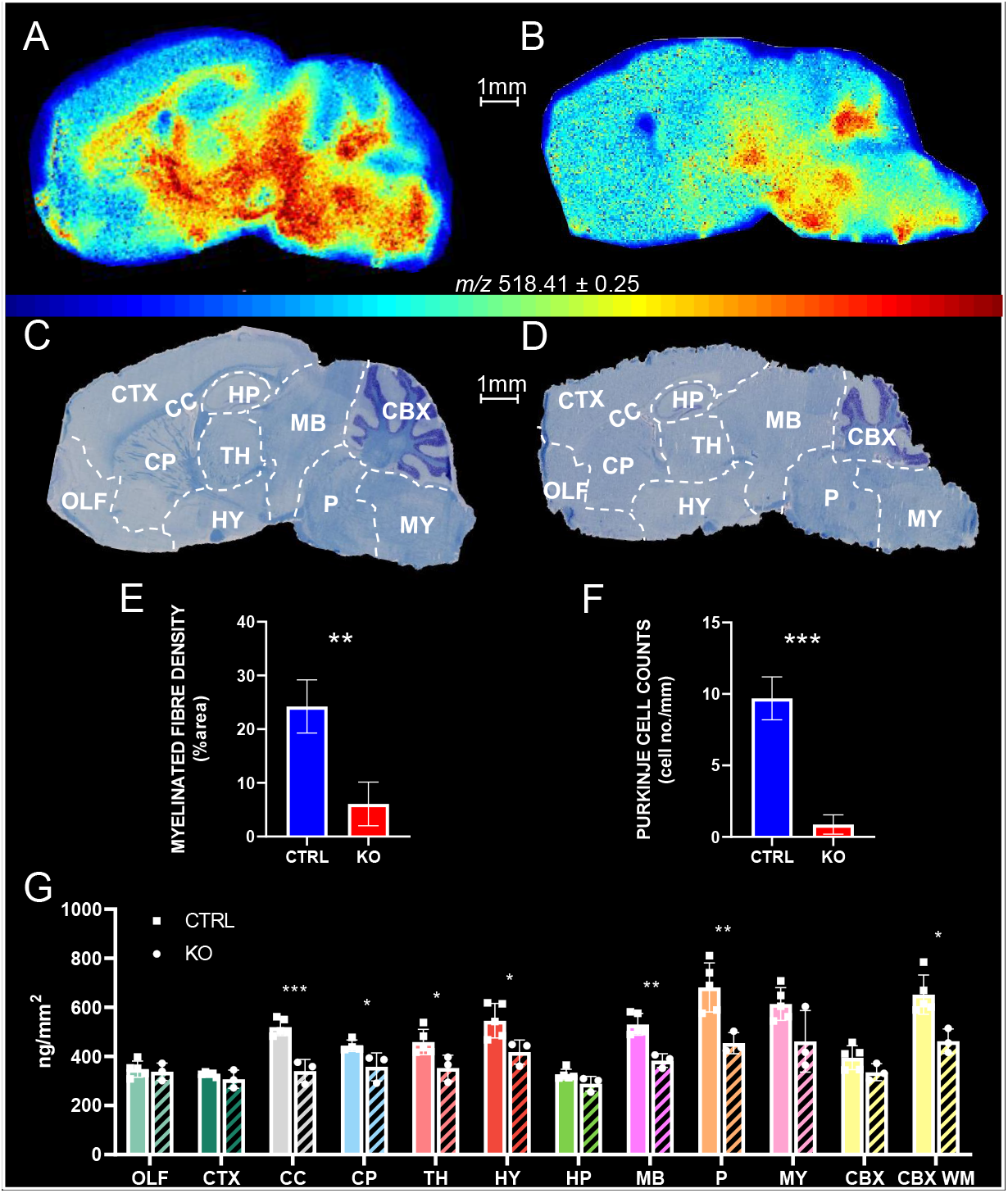
Histology-matched quantitative MALDI-MSI data displaying cholesterol distribution and quantification in brain tissue sagittal sections of WT and *Npc1*^-/-^ mice. (A, B) MALDI-MSI showing cholesterol distribution in WT (A), and in *Npc1*^-/-^ (B) mouse brain tissue. The cholesterol signal was normalized to the signal of sprayed-on [^2^H_7_]cholesterol. Data acquired using vacuum-MALDI-TOF MS is shown using a “jet” scale. Scale bar 1 mm. Pixel size 50 µm. (C, D) LFB/CV staining for myelin of sagittal mouse brain sections adjacent to sections undergoing MSI. Major anatomical structures were identified by comparison with the corresponding Reference Atlas of the adult mouse brain provided by the Allen Institute of Brain Science (Lein *et al*., 2007) and are outlined with dashed lines. Abbreviations are the same as in Figure 3. (E) Myelinated fibre density in the CP of WT and of *Npc1*^-/-^ mice as assessed in LFB/CV stained sagittal sections. (F) Purkinje cells counts in the CBX of WT and *Npc1*^-/-^ mice as assessed in LFB/CV stained sagittal sections. (G) Areal density (ng/mm^2^) of cholesterol in brain regions from five WT and three *Npc1*^-/-^ mice, averaged over the biological replicates (see *Statistics* section. WT, n = 5, 3 sections per mouse for a total of 15 measurements, *Npc1*^-/-^, n = 3, 3 sections per mouse for a total of 9 measurements). Significance levels from a Shapiro-Wilk test for normality followed by an unpaired *t*-test are indicated for those brain regions showing significant differences. *P < 0.05, **P < 0.01, ***P < 0.001. Average values for WT and *Npc1*^-/-^ mice groups are given by separate histogram bars. The number of dots within each bar indicates the number of mice analysed for each group. Each dot within the bars correspond to individual average for each mouse. The height of each bar represents the mean of the region average for each mouse group (▪CTRL, or •KO). The error bars indicate the SD of all the replicates (sections) per mouse.

As with the WT mouse, cholesterol areal densities in the *Npc1*^-/-^ mouse were determined against a known density of sprayed-on internal standard. The quantitative data reported in Figure 6G and in Table 1 indicate that cholesterol abundance follows the pattern: cerebellar white matter (462.1 ± 58.7 ng/mm^2^) ≈ medulla (461.6 ± 111.3 ng/mm^2^) ≈ pons (454.2 ± 49.5 ng/mm^2^) > hypothalamus (417.7 ± ng/mm^2^) > midbrain (383.1 ± 32.0 ng/mm^2^) > caudate-putamen (357.5 ± 50.7 ng/mm^2^) ≈ thalamus (352.6 ± 51.9 ng/mm^2^) > corpus callosum (340.8 ± 44.6 ng/mm^2^) ≈ olfactory traits (337.5 ± 35.2 ng/mm^2^) ≈ cerebellum (335.8 ± 40.0 ng/mm^2^) > cortex (307.4 ± 36.9 ng/mm^2^) > hippocampus (288.9 ± 36.5 ng/mm^2^).

Table 1 reports cholesterol areal density values for the *Npc1*^-/-^ mouse as measured by EADSA-MSI and % differences compared with the WT animal. The regions showing highest reduction of cholesterol in the *Npc1*^-/-^ mouse compared to the WT are the corpus callosum (34.4%), pons (33.4%), cerebellar white matter (29.1%), midbrain (27.8%), medulla (24.8%), hypothalamus (23.5%), thalamus (23.2%) and caudate-putamen (13.6%). All these differences, except for the medulla, are significant (p-values are indicated in Figure 6G).

A comparison of histological as well as MSI data in WT and *Npc1*^-/-^ mouse brain reveals structural differences that can be correlated with compositional changes of cholesterol distribution and abundance. The most striking difference is in the CC. Figure 6C shows that the CC in the WT mouse is heavily myelinated and highlighted by the LFB dye. On the contrary, Figure 6D shows that the CC is apparently non-myelinated in the *Npc1*^-/-^ brain with the LFB stain showing this structure as mostly white. This correlates well with our MSI data where cholesterol areal density is significantly higher in the WT as compared to the *Npc1*^-/-^ CC (Figure 6G, ***p-value < 0.001).

Other than in the CC, the significantly higher cholesterol areal density as determined by MSI in the caudate-putamen region, and in the cerebellar white matter of the WT mouse as compared to the *Npc1*^-/-^ mouse (*p-values < 0.05, Figure 6G), also relates to known histological markers (German *et al*., 2002, German *et al*., 2001). This prompted us to further analyse histological data by assessing the percentage of myelinated fibres in the caudate-putamen (Figures 6E and Supplemental S5A), and the number of Purkinje cells in the cerebellar GM (Figures 6F and Supplemental S5B) of these mice.

In the caudate-putamen, myelinated fibre density assessed in LFB/CV stained sections was found to be significantly reduced in the *Npc1*^-/-^ mouse as compared to WT (Figures 6E and Supplemental S5A, **p-value < 0.01), agreeing with MSI measurement of cholesterol areal density in the same brain region (Figure 6G, *p-value < 0.05).

Focusing on the cerebellum, the MSI quantitative measurements show a statistically significant difference when cerebellar WM is considered (Figure 6G, *p-value < 0.05). This data is in agreement with a previous MSI study (Tobias *et al*., 2018), similarly showing a reduced cholesterol signal intensity, as normalized by total ion current (TIC), in the cerebellum of the same *Npc1*^-/-^ mouse when compared to WT. Interestingly, there is significant reduction of the number of Purkinje cells in the GM of the *Npc1*^*-/-*^ cerebellum (Figures 6F and Supplemental S5B, ***p-value < 0.001). Loss of Purkinje cells is a well know phenotypic marker of NPC patients (Gilbert *et al*., 1981) and animal models (Higashi *et al*., 1993). These neuronal cells have their cell bodies residing in the cerebellar GM, but their myelinated axons establish postsynaptic connections with cerebellar deep nuclei in the WM (Goodlett & Mittleman, 2017). Therefore, a reduction in the cholesterol content of the cerebellar WM of the *Npc1*^-/-^ mouse could be explained in part by the loss of Purkinje cell efferent and afferent connections.

A significant reduction in the cholesterol areal density of hypothalamus, midbrain, and pons in the *Npc1*^-/-^ mouse compared to WT was revealed by MSI (Figure 6G, *p-value < 0.05 and ** p-value < 0.01). Observation of the histological staining shown in Figure 6C and 6D illustrates reduced LFB stain density in these same areas of the *Npc1*^-/-^ mouse, correlating with MSI heatmaps of cholesterol in Figures 6A and 6B, respectively.

### EADSA-MSI with Multiple Ionisation Modes and Analysers

We assessed the robustness of our EADSA-MSI method on different mass spectrometers having different sources (vacuum-MALDI, AP-MALDI and DESI) and analysers (Q-TOF with and without ion mobility, linear TOF, Orbitrap). Our data is consistent as shown for the cholesterol distribution in adult mouse in Figures 1C, 5F, 6A, Supplemental S2A all generated by vacuum-MALDI-TOF (Bruker ultrafleXtreme), in Supplemental Figure S4A generated by MALDI-Q-IM-TOF (Waters Synapt G2), in Supplemental Figure S4C generated by DESI-Q-TOF (Waters Synapt G2), and in Figure 4A generated by AP-MALDI-Orbitrap (MassTech and ThermoFisher Scientific). Supplemental Figures S4A and S4C represents sagittal sections of different WT mice and were taken on different planes separated by about 300 µm. Supplemental Figure S4A is matched with the correspondent reference section from the Allen Mouse Brain Atlas (Lein *et al*., 2007) in Figure S4B, and Figure S4C is matched with the appropriate atlas section in Figure S4D. Altogether these data illustrate the reproducibility and robustness of our EADSA-MSI methodology.

## Conclusion

The EADSA-MSI method presented provides a tool for the quantitative imaging of cholesterol in histological mouse brain tissue sections. On-tissue EADSA was successfully employed to improve the analytical power of MSI toward sterols, allowing quantitative mapping of cholesterol at pixel sizes down to 30 µm. Different MS platforms were utilized including vacuum-MALDI-TOF, vacuum-MALDI-Q-IM-TOF, AP-MALDI-Orbitrap and DESI-Q-TOF, demonstrating the robustness of the method towards different ionization sources, analysers and detectors. With atmospheric pressure ionisation-MSI (AP-MALDI-Orbitrap and DESI-Q-TOF) the method allowed detection of other lipid classes (phospholipids) in parallel to derivatised sterols, thereby extending the reach of the methodology to the characterization of diverse lipid markers simultaneously. MSI is a rapidly advancing technology that can reach cellular resolution (Soltwisch *et al*., 2020), thereby providing information not only on structural changes but also on changes happening at the level of different cell populations. Bridging MS-based lipidomics with histopathology will allow the correlation of high sensitivity quantitative molecular information with anatomical location, opening a further window for the entry of MSI into clinical chemistry. The EADSA-MSI method described here for imaging of cholesterol directly on tissue can be easily applied to a number of scientific fields including neuroscience, pharmacology, biochemistry and pathology. Particularly, its application to the study of diseases such as Alzheimer’s and multiple sclerosis has the potential to unveil the role of cholesterol in these important neuropathologies.

## Material and Methods

The aim of the study was to develop an MSI method suitable to map the distribution and to determine the concentration of cholesterol in different anatomical regions of mouse brain.

### Chemicals and Reagents

HPLC grade methanol, propan-2-ol, acetonitrile, ethanol, xylene, industrial methylated spirit and water were from Fisher Scientific (Loughborough, UK). Glacial acetic acid was purchased from VWR (Lutterworth, UK). [25,26,26,26,27,27,27-^2^H_7_]Cholesterol was from Avanti Polar Lipids (Alabaster, AL). Cholesterol oxidase from *Streptomyces sp*., potassium dihydrogen phosphate, Luxol Fast Blue (LFB), Cresyl Violet (CV), DPX mountant, paraformaldehyde (PFA), and α-cyano-4-hydroxycinnamic acid (CHCA) were from Sigma-Aldrich, now Merck (Dorset, UK). GP-hydrazine was from TCI Europe (Zwijndrecht, Belgium). Lithium carbonate was from Acros Organic, now Thermo Fisher Scientific (Geel, Belgium).

### Experimental models

For the present study WT and *Npc1*^-/-^ mice were employed. Experiments were performed in accordance with University of Illinois at Chicago IACUC approved protocols. Balb/c *npc*^nih^ (*Npc1*^*+/-*^) mice were obtained from Jackson Laboratories (RRID:IMSR JAX:003092) and a breeding colony was maintained. Genotype was confirmed by PCR as previously reported (Tobias *et al*., 2018). At ten weeks of age, WT (*Npc1*^*+/+*^) and the null mutant (*Npc1*^*-/-*^) mice were euthanized via CO_2_ asphyxiation followed by decapitation. All mice were males. In all cases, whole brain was removed and immediately frozen in dry ice to maintain spatial integrity and stored at -80°C. Details of the phenotypically normal 1-day-old newborn mouse can be found in Supplemental Methods.

### Tissue Sectioning

Fresh frozen brain tissue, mounted on and only partially embedded in OCT (Optimal Cutting Temperature) compound, was cryo-sectioned using a Leica Cryostat CM1900 (Leica Microsystems, Milton Keynes, UK) at a chamber temperature of -18°C into 10 μm-thick sections which were thaw-mounted onto optical microscope slides for histology or onto indium tin oxide (ITO) coated glass slides for MSI, and stored at -80°C until use. ITO coated glass slides (8 - 12 Ohm/Sq) for MSI were from Diamond Coatings (Halesowen, UK). Three sections were mounted on each glass slide, each section was separated by 100 µm from the adjacent section, i.e. the nine sections in between were placed on other consecutive slides.

### Histology

Tissue sections adjacent to sections analysed by MSI were thawed, fixed in PFA to preserve anatomy, and subjected to LFB histology with cresyl violet as the counterstain, essentially as described in (Kluver & Barrera, 1953) and in Supplemental Methods. Finally, a cover slip was placed over the specimens with DPX permanent mountant. Histological data was analysed by QuPath (Bankhead *et al*., 2017) and ImageJ (Schneider *et al*., 2012) following whole section digitisation at 400x magnification using a Zeiss AxioScanner.

### Region of Interest Analysis and Quantitative Morphometry

Quantitative analysis of histological data was carried out as follows. To assess fibre myelination, the caudate-putamen ROI was outlined on the digitised images with Qupath (Supplemental Figure S5A). Images of defined ROI were cropped and converted into an 8-bit format with ImageJ to mark and measure specific areas. The threshold was adjusted to exclude cell nuclei and automatically outline WM areas exclusively, and total WM area was measured per section. These data were used to calculate the percentage of myelinated fibres in the selected ROI (Figure 6E). Cerebellar area and length of the Purkinje cell layer were also defined with Qupath (Supplemental Figure S5B) and Purkinje cells were manually counted to determine cell linear density (Figure 6F). Statistical analysis was carried out on GraphPad 8.1. A Shapiro-Wilk test was employed to confirm normality followed by an unpaired *t*-test to assess statistical significance of measured differences.

### Stereology

Stereological methods were employed to identify ROI within mouse brain sagittal tissue sections. The defined ROI where employed to analyse both the MSI and the histology data. For determination of ROI we referred to the Allen Mouse Brain Atlas (sagittal sections, P56, https://atlas.brain-map.org/atlas?atlas=2) (Lein *et al*., 2007). Firstly, the mouse brain was sectioned through a sagittal plane. Then stained sagittal sections were matched with images in the corresponding reference atlas (Allen Institute for Brain Science). The shape of the fibres tracts as detected by LFB was used to identify the correct atlas reference image having a comparable distance from the midline. Then the LFB/CV stained sections and the MSI heat maps were overlaid together with the appropriate reference atlas images, and anatomical regions identified were outlined with dashed lines defining ROI (Figures 3A, 6C-D, and Supplemental S6).

For MALDI-MSI data analysis of the adult mouse, three sagittal sections at about 1.3 mm ± 300 µm distance from the midline were employed. The three chosen sections were taken at sagittal planes separated by 100 µm. For histology data analysis we employed sections adjacent to MSI on the left and on the right for a total of 6 sections per mouse that covered the area 1.3 mm ± 400 µm distant from the midline. The AP-MALDI data in Figure 4 was an exception and obtained on a more peripheral section (about 3 mm from the midline). The mouse brain atlas images (credit: Allen Institute) employed as reference were: Adult Mouse, P56, Sagittal, Images 13-16 of 21 ids=100883818, 100883869, 100883867, 100883888 (Lein *et al*., 2007).

For MALDI-MSI data analysis of the newborn mouse, four consecutive sagittal sections at about 0.6 mm ± 100 µm from the midline were employed. For histology, two sections adjacent on the far left and on the far right of the series of four used for MSI were employed. The newborn mouse brain atlas image (credit: Allen Institute) employed as reference was Developing Mouse, E18.5, Sagittal, Image 16 of 19 id= 100740373 (Lein *et al*., 2007).

### Deposition of Internal Standard and On-Tissue EADSA

This was performed as described by (Yutuc *et al*., 2020) with minor modifications. Frozen mouse brain sections mounted on a ITO coated glass slide were transferred under dry ice from the -80°C freezer to a vacuum desiccator in which the pressure was reduced to 0.1 bar allowing the tissue to dry for 15 min. Afterwards, [^2^H_7_]cholesterol (200 ng/μL in ethanol) was sprayed from a SunCollect automated pneumatic sprayer (SunChrom, Friedrichsdorf, Germany supplied by KR Analytical Ltd, Cheshire, UK) at a flow rate of 20 μL/min at a linear velocity of 900 mm/min with 2 mm line distance and height of 30 mm from the section in a series of 18 layers. The resulting density of the deuterated standard was 40 ng/mm^2^ (see below). The sprayer was thoroughly flushed with about 2 mL of methanol after which cholesterol oxidase (0.264 units/mL in 100 μM KH_2_PO_4_ pH 7) was sprayed for 18 layers. The first layer was applied at 10 μL/min, the second at 15 μL/min, then all the subsequent layers at 20 μL/min to give an enzyme density of 0.05 munits/mm^2^. Thereafter, the enzyme-coated slide was placed on a custom-made dry PTFE bed in a glass staining jar (11 cm x 11 cm x 7.5 cm) above 30 mL of warm water (37°C), then incubated at 37°C for 1 hr. Subsequently, the slide was removed, and the tissue was dried in a vacuum desiccator for 15 min as described above. GP (5 mg/mL in 70% methanol, 5% acetic acid) was sprayed on the dried slide with the same spray parameters as used for spraying of cholesterol oxidase. The resulting GP density was 1.00 μg/mm^2^. The slide was then placed in the custom-made humidity chamber as above containing 10 mL of pre-warmed (37°C) 50% methanol, 5% acetic acid and incubated at 37°C for 1 hr. The slide was removed and dried in a vacuum desiccator as above, then stored in a cold room (4°C) until MSI analyses. DESI-MSI experiments were performed without any further pre-treatment. For MALDI-MSI, on the next day the desiccator was allowed to reach room temperature, and then the slide was removed and sprayed with CHCA MALDI matrix. CHCA was sprayed from a HTX TM-Sprayer (HTX Technologies, NC, USA) at 5 mg/mL in water:propan-2-ol:acetonitrile (3:4:3, v:v:v) at a flow rate of 80 μL/min and a linear velocity of 1200 mm/min, with 2 mm line distance and a criss-cross deposition method which alternates vertical and horizontal passes, for a total of 8, with an offset of 1 mm, resulting in a matrix density of 1.33 μg/mm^2^. The sprayer nozzle was heated at 70°C to enhance solvent evaporation rate.

### MSI

Following EADSA treatment, tissues sections were analysed using different mass spectrometers. Optimized instrumental parameters are described below.

### Vacuum MALDI-TOF-MSI

Experiments were carried out on an ultrafleXtreme MALDI TOF/TOF mass spectrometer (Bruker Daltonics, Bremen, Germany) equipped with a Smartbeam™ Nd:YAG laser emitting at 355 nm (2 kHz) and operated in the reflectron mode and positive polarity. Each mass spectrum was automatically acquired using the auto-execute method in FlexControl (Bruker) software in the range of *m/z* 400 – 1000. Pixel size was set at 50 μm using flexImaging 4.1 software (Bruker), setting laser focus to “small”. Laser power was tuned to optimise signal to noise (S/N) ratio without distortions of the baseline at 80% of maximum with Global Offset at 5%, Attenuator Offset at 40%, Attenuator Range at 40%. These parameters result in a diameter of the laser spot of about 50 μm, according to factory specifications and as verified by visual inspection with the instrument camera. Extraction voltages were as follows: IS1 20.00 – 19.92 kV, IS2 17.90 – 17.82 kV, Lens 8.50 – 8.53 kV, Rfl 21.10 – 20.99 kV, Rfl2 10.95 – 10.89 kV. Reflector gain was set at 3.0X. Pulsed Ion Extraction was timed at 160 ns. A gated ion suppression was applied up to *m/z* 375. Each raster was sampled with 200 shots in 5 steps for a total of 1000 shots per raster. Total acquisition time was typically about 11.5 hr for a total of ∼27000 positions and a file size of ∼24000 MB. The MALDI instrument was calibrated using a mixture of phosphatidylcholine and lysophosphatidylcholine (Avanti Polar Lipids) of known composition and having masses in the range of interest. After measurement, imaging spectra were re-calibrated using the batch process in flexAnalysis. On-tissue, mass accuracy was typically within ∼100 ppm of the theoretical mass. Data were analysed and visualized using flexImaging 3.0 (Bruker), and SCiLS Lab 2014b (SCiLS, Bremen, Germany) without any processing step. Data were visualized using normalization to [^2^H_7_]cholesterol at *m/z* 525.5. Mass selection windows for ion of interests were chosen with a width of ± 0.25 Da in flexImaging 3.0 and of ± 0.125% in SCiLS Lab 2014b. A mass resolution (full width at half-maximum) of ∼M/ΔM 20000 was typically achieved in a single pixel. An optical image of each tissue section was acquired prior to the MS acquisition by means of a flatbed scanner.

### AP-MALDI-MSI

MSI experiments were carried out in the positive-ion mode with an Orbitrap Elite™ Hybrid Ion Trap-Orbitrap Mass Spectrometer (ThermoFisher Scientific) coupled with an AP-MALDI UHR Source (MassTech, Maryland USA, supplied by KR Analytical Ltd) equipped with a Nd:YAG laser emitting at 355 nm. Full scan mode (MS^1^) imaging analysis was performed with *m/z* measurement in the Orbitrap over the *m/z* range of 400 - 1200 at 60,000 resolution (FWHM at *m/z* 400), MALDI laser energy was set at 45% of maximum and frequency was 1.5 kHz. Data were acquired in Constant Speed Raster (CSR) mode at a scan speed of 2.8 mm/min and a pixel size of 30 µm. Capillary temperature was 450°C, voltage was 3 kV, injection time was 400 ms, and a lock mass for ^2^[H_7_]cholesterol at *m/z* 525.4544 was employed. The acquisition of one mouse brain tissue section was achieved in about 15 hours for a total of about 90,000 positions and a file size of about 6 GB. In MS^3^ experiments, the MALDI laser energy was set at 14% and frequency was 1.5 kHz. Data were acquired in CSR mode at a scan speed of 3 mm/min and a pixel size of 40 µm. Capillary temperature was 350°C, voltage was 3.5 kV, and injection time was 300 ms. In the linear ion trap (LIT), precursor ions were isolated and fragmented with an isolation width of 2 and an arbitrary collision induced dissociation (CID) energy of 35%. The most intense fragment ion produced in MS^2^ was selected with an isolation width of 2 and fragmented with a CID energy of 40% to produce an MS^3^ spectrum. MS^3^ fragmentation spectra of cholesterol and ^2^[H_7_]cholesterol were acquired in each pixel. The MS^3^ transition for cholesterol was 518.4 **→** 439.4**→**. The MS^3^ transition for ^2^[H_7_]cholesterol was 525.4 **→** 446.4 **→**.

Data were analysed and visualized using ImageQuest (ThermoFisher Scientific). Alternatively, after exporting the file into an imzml format, data were analysed by MSiReader (Bokhart *et al*., 2018), and SCiLS Lab 2014b (SCiLS, Bremen, Germany) without any processing step. MS^1^ data were normalized to the isotope-labelled [^2^H_7_]cholesterol at *m/z* 525.454. The ions of interests were extracted with a width of 7 mDa for the full scan experiment and 0.3 Da for the MS^3^ experiment.

### MALDI-Q-IM-TOF MSI

Experiments were carried out on two Synapt G2-Si instruments (Waters, Wilmslow, UK) exploiting Waters HDI 1.4 software. The same software was employed for image visualization. Images were generated from spectra acquired in the positive ion mode in the *m/z* range 400 - 1000. The laser frequency was 1 kHz, and power was kept at 100 arbitrary units. Scan time was 0.5 sec and pixel size was 50 µm. IMS cell Wave Velocity was from 1000 to 300 m/s, and Transfer Wave Velocity was 281 m/s. In all experiments the cholesterol signal was measured to better than 5 ppm mass accuracy.

### DESI-Q-IM-TOF MSI

Experiments were carried out on a Synapt G2-Si (Waters, Wilmslow, UK) using Waters HDI 1.4 software. The same software was employed for image visualization. Spectra were acquired in the positive ion mode in the *m/z* range 100 - 1200, with a needle voltage of 4.5 kV. The DESI solvent flow rate was 1.25 µL/min. Scan time was 0.25 s, and pixel size was 25 µm. IMS cell Wave Velocity was from 1000 to 300 m/s, and Transfer Wave Velocity was 281 m/s. In all experiments the cholesterol signal was measured with a mass accuracy better than 8 ppm.

### Quantification

To achieve reliable quantitative measurements, known amounts of [^2^H_7_]cholesterol were sprayed on-tissue prior to the EADSA process. This procedure corrects for variation in extraction efficiency, background matrix effect, and ultimately MS response. The linearity of the on-tissue response of sprayed-on [^2^H_7_]cholesterol verses endogenous cholesterol was determined by spraying eight consecutive tissue sections with [^2^H_7_]cholesterol at varying densities (endogenous cholesterol areal density is assumed to be constant for a given ROI across the consecutive slices, Supplemental Figure S2A). Examples of calibration curves obtained on whole-brain sections and considering the cerebellum only as a ROI are shown in Supplemental Figures S2B and S2C, respectively. Quantification was made from [M]^+^ ion signal intensities, averaged in each brain region. Brain regions of interest were defined according to Allen Mouse Brain Atlas. During cryosectioning of the sagittal brain sections, the distance from the brain midline was controlled and kept at about 1.3 mm ± 300 µm for the adult mouse (except for experiments made with AP-MALDI-MSI), and at about 0.6 mm ± 100 µm for the newborn mouse. The areal density of cholesterol in defined regions of interest was calculated by correlating signal intensity to that of known density of [^2^H_7_]cholesterol sprayed on-tissue. All quantitative measurements were made employing SCiLS Lab MVS (2019c Core, SCilS, Germany).

### Statistics

Statistical analysis was applied to the quantitative assessment of cholesterol, myelinated fibre density, and specific cell counts, in defined brain regions of WT and *Npc1*^*-/-*^ mouse brain. Five WT and three *Npc1*^*-/-*^ brains were employed, analysing three or more sections for each mouse. To determine statistical difference in cholesterol areal density between defined regions of interest in five adult WT mice, two-way ANOVA was performed with cholesterol areal density as dependent variable and mouse and brain region as factors. The interaction between mouse and brain region was used as error variance. The residuals representing the interaction deviations were approximately normally distributed. Tukey’s multiple comparisons test was used to identify significant differences between brain regions.

To determine statistical differences in cholesterol areal density in defined regions of interest between WT and *Npc1*^-/-^ mouse brain a Shapiro-Wilk test for normality was performed, followed by an unpaired *t*-test for significance. The analyses were performed using GraphPad Prism 8.2.1 software (GraphPad Software, Inc, CA, USA). A *P*-value of less than 0.05 was considered statistically significant. *P*<0.05, *; *P*<0.01, **; *P*<0.001, ***. All whiskers on bar graphs represent 1 standard deviation. Note, one of the 5 control mice was not considered in the calculation of the average cholesterol areal density for the caudate-putamen as it was not sectioned on an equivalent anatomical plane.

### Calculations of areal densities of internal standard, cholesterol oxidase and GP-hydrazine

To calculate areal densities of [^2^H_7_]cholesterol, cholesterol oxidase and GP-hydrazine we employed the following equation:

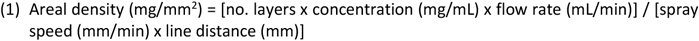

[^2^H_7_]Cholesterol in ethanol (200 ng/μL) was sprayed on-tissue using a SunCollect automated pneumatic sprayer at a flow rate of 0.02 mL/min, a spray speed of 900 mm/min, a line distance of 2 mm, for 18 layers. Using (1), the areal density on brain of [^2^H_7_]cholesterol was calculated to be 40 ng/mm^2^.

The cholesterol oxidase activity was 0.264 U/mL in the sprayed solution, using eq.1 and the same spray parameters as for the internal standards, this translates to an areal density of 0.05 mU/mm^2^.

The concentration of the chloride salt of GP sprayed on tissue was 5 mg/mL, using the same spray parameters as above this translated to an areal density of 1.00 μg/mm^2^ (5.1 nmol/mm^2^).

## Acknowledgements

This work was supported by UKRI Biotechnology and Biological Sciences Research Council (BBSRC, grant numbers BB/N015932/1 to WJG/YW/OWH/MRC/JN, BB/L001942/1 to YW). RA holds a MSCA-COFUND Sêr Cymru Fellowship supported by the Welsh Government and the European Regional Development Fund. Work at the Nebraska Medical Center was supported by the NIH NIMH MH110636 (National Institute of Health National Institute of Mental Health). Work at the University of Illinois at Chicago was supported by the Ara Parseghian Medical Research Fund and the Department of Chemistry, College of Liberal Arts and Sciences. Dr Ruth Andrew (University of Edinburgh) is thanked for advice on performing on-tissue derivatisation. We acknowledge Dr Rosalind John for kindly providing the CD1 mice and Bridget Allen for brain dissection (Cardiff University). Dr Tina Angerer (Luxembourg Institute of Science and Technology) is thanked for helpful discussions. We are grateful to Professor David O.F. Skibinsky (Swansea University) for providing statistical advice.

