## Supplemental Methods for "Visualising Cholesterol in Brain by On-Tissue Derivatisation and Quantitative Mass Spectrometry Imaging"

* Corresponding author: William J Griffiths

ORCIDs: RA, 0000-0001-5136-5921; EY, 0000-0001-9971-1950; MFW, 0000-0003-4107-5941; JN, 0000-0003-4848-7391; ZK, 0000-0002-8690-4507; SMC, 0000-0002-3541-3361; OWH, 0000-0003-2157-9157; MRC, 0000-0002-0798-831X; YW, 0000-0002-3063-3066; WJG, 0000-0002-4129-6616.

**Keywords**

Mass Spectrometry Imaging; Quantification; MALDI; Brain; Derivatisation; Cholesterol; Myelin, Sterol; Development; Niemann-Pick disease;

**Supplemental Methods**

The aim of the study was to develop an MSI method suitable to map the distribution and to determine the concentration of cholesterol in different anatomical regions of the mouse brain.

*Experimental models*

To study cholesterol distribution during mouse development, the phenotypically normal *Dhcr7*$^{T93M/+}$ was used (Correa-Cerro *et al.*, 2006). In the present study one-day-old newborn animals (P0) were employed. All mice were housed under a 12 hr light-dark cycle at constant temperature (25 °C) and humidity with *ad libitum* access to food (Teklad LM-485 Mouse/Rat Irradiated Diet 7912) and water at the University of Nebraska Medical Center. Newborn mice were used for the study. All procedures were performed in accordance with the Guide for the Humane Use and Care of Laboratory Animals. The use of mice in this study was approved by the Institutional Animal Care and Use Committee of University of Nebraska Medical Center. Note our initial aim was to also image brain from the newborn *Dhcr7*$^{T93M/\Delta3\text{-}5}$ mouse, however, brain from this animal proved particularly difficult to section, and will require further investigation.

Adult mice for exploratory experiments were kindly provided by Dr Rosalind John and brain dissected by Bridget Allen (Cardiff University). Mice were CD1, females, 7 months old. Mice were euthanized by dislocation of the neck (schedule 1) in accordance with institutional animal care guidelines and brains dissected immediately post-mortem and snap-frozen in liquid nitrogen. Mice were housed in a conventional unit on a 12-hr light–dark cycle with lights coming on at 06:00, with a temperature range of 21 ± 2°C, and with free access to tap water and standard chow. All procedures were conducted in accordance with the requirements of the UK Animals (Scientific Procedures) Act 1986, under the remit of Home Office licence (BA) with additional ethical approval at Cardiff University.

*Histology*

Luxol Fast Blue and Cresyl Violet staining was performed according to (Kluver & Barrera, 1953) on tissue sections adjacent to sections analysed by MSI. Snap frozen sections previously stored at -80°C, were allowed dry at room temperature for 20 min, and then incubated with 4% PFA for 1 hr (~100 µL on each section). Afterwards, the sections were dehydrated in 70% then 95% ethanol, for 2 min each. Subsequently, sections were incubated overnight (< 16 hr) in a sealed jar at 40°C in 0.1% LFB in 95% ethanol. The LFB solution was filtered and preheated before incubation. Afterwards, excess stain was removed by gently washing the sections in a saturated solution of lithium carbonate until WM and GM could be distinguished. The LFB stained slides were counterstained with 0.1% Cresyl Violet acetate solution preheated at 65°C for 30 min and then quickly washed in water and dehydrated in 95% then absolute ethanol (1-2 seconds each). Afterwards the slides were cleared in xylene overnight and dried at room temperature for a few seconds.

Supplemental Figure S1. Fragmentation pattern of GP-derivatised cholesterol upon $MS^n$ analysis showing the chemical structures of the major diagnostic fragment ions.

Supplemental Figure S2. Generation of calibration plots for endogenous cholesterol against sprayed-on $[^2H_7]$cholesterol on mouse brain tissue. To determine absolute quantities of cholesterol, calibration curves were constructed by spraying a solution of $[^2H_7]$cholesterol at varying concentrations onto consecutive mouse brain tissue sections. (A) From left to right and top to bottom, MSI depicting cholesterol normalized to the peak of $[^2H_7]$cholesterol sprayed on-tissue at decreasing concentration of $[^2H_7]$cholesterol. (B) Calibration curve generated using the entire section. (C) Calibration curve generated using the cerebellar area outlined in red. Data normalized to sprayed-on $[^2H_7]$cholesterol are shown using a "jet" scale over a single range. Scale bar = 1 mm, spatial resolution 50 µm.

Supplemental Figure S3. (A) Gross anatomy of the corpus callosum. The corpus callosum is approximately 7 mm in length and is C-shaped in a gentle upwardly convex arch. It is divided into four parts (from anterior to posterior): rostrum (continuous with the lamina terminalis), genu, trunk/body, splenium. Body and splenium are connected by the thinnest isthmus. (B) Reproducibility of the morphology of the Corpus Callosum in the five WT mice analysed, one section per mouse is shown. An example of the ROI for CC structure is also shown. (C) Areal density ($ng/mm^2$) of cholesterol in brain regions from five WT mice, averaged over the sections (n = 5 mice, 3 sections per mouse for a total of 15 measurements). Average values for CC regions in the WT mice group are given by separate histogram bars. The height of each bar represents the mean of the region average in the WT mice group. The error bars indicate the SD of all the replicates (sections) per mouse.

Supplemental Figure S4: MSI depicting cholesterol in WT mouse brain tissue obtained by on-tissue EADSA derivatisation and subsequent MS analysis together with corresponding reference atlas sagittal sections. (A) MALDI-IM-MSI of adult mouse brain cholesterol and (B) reference atlas section corresponding to the same sagittal plane. Image credit Allen Institute: Mouse, P56, Sagittal, Image 11 of 21 id= 100884129 (Lein *et al.*, 2007). (C) DESI-IM-MSI of cholesterol of adult WT mouse brain and (D) corresponding reference atlas sagittal section. Image credit Allen Institute for Brain Science: Mouse, P56, Sagittal, Image 9 of 21 id= 100883813 (Lein *et al.*, 2007). (E) DESI-IM-MSI of cholesterol of 1-day-old WT mouse brain and (F) corresponding reference atlas sagittal section. Image credit Allen Institute for Brain Science: Developing Mouse, E18.5, Image 16 of 19 id=100740373 (Lein *et al.*, 2007). For MSI experiments: isolation window width 7 mmu, spatial resolution 50 µm (A) and 45 µm (B-C); data analysed by Mass Lynx (Waters) visualised on a BPROY scale. All images data are normalised to sprayed-on $[^2H_7]$cholesterol.

Supplemental Figure S5: LFB/CV histological staining of WT (left panels) and *Npc1*$^{-/-}$ mouse brain (right panels). (A) Brain region of the caudate-putamen showing myelinated fibres in the WT but absent from the *Npc1*$^{-/-}$ mouse. In the enlargements a single fibre can be seen in the WT. (B) Cerebella region showing (with Qu Path annotation for Purkinje cells) the difference between WT and *Npc1*$^{-/-}$ in the number Purkinje cells. In the enlargements a stream of eight Purkinje cell is shown.

Supplemental Figure S6: Example of definition of ROI outlines in SCiLS Lab software (SCiLS, Bremen, Germany).

Supplemental Table S1: Cholesterol areal density values in defined brain areas of two WT newborn mice as quantified by MALDI-MSI. Four replicate sections per each mouse were employed. Averages ± standard deviation across all replicates are reported.

| Brain area | Cholesterol (ng/mm$^2$) |
|---|---|
| pons | 125.1 ± 15.3 |
| diencephalon | 160.5 ± 24.0 |
| mid brain | 171.9 ± 14.1 |
| prepontine hindbrain | 175.6 ± 23.2 |
| pontine hindbrain | 193.4 ± 28.4 |
| pontomedullary and medullary hindbrain | 180.3 ± 25.6 |
| peduncular hypothalamus | 162.2 ± 19.8 |
| telencephalic vesicle | 111.7 ± 13.9 |
