## Supplemental Figure for "Visualising Cholesterol in Brain by On-Tissue Derivatisation and Quantitative Mass Spectrometry Imaging"

Figure S1

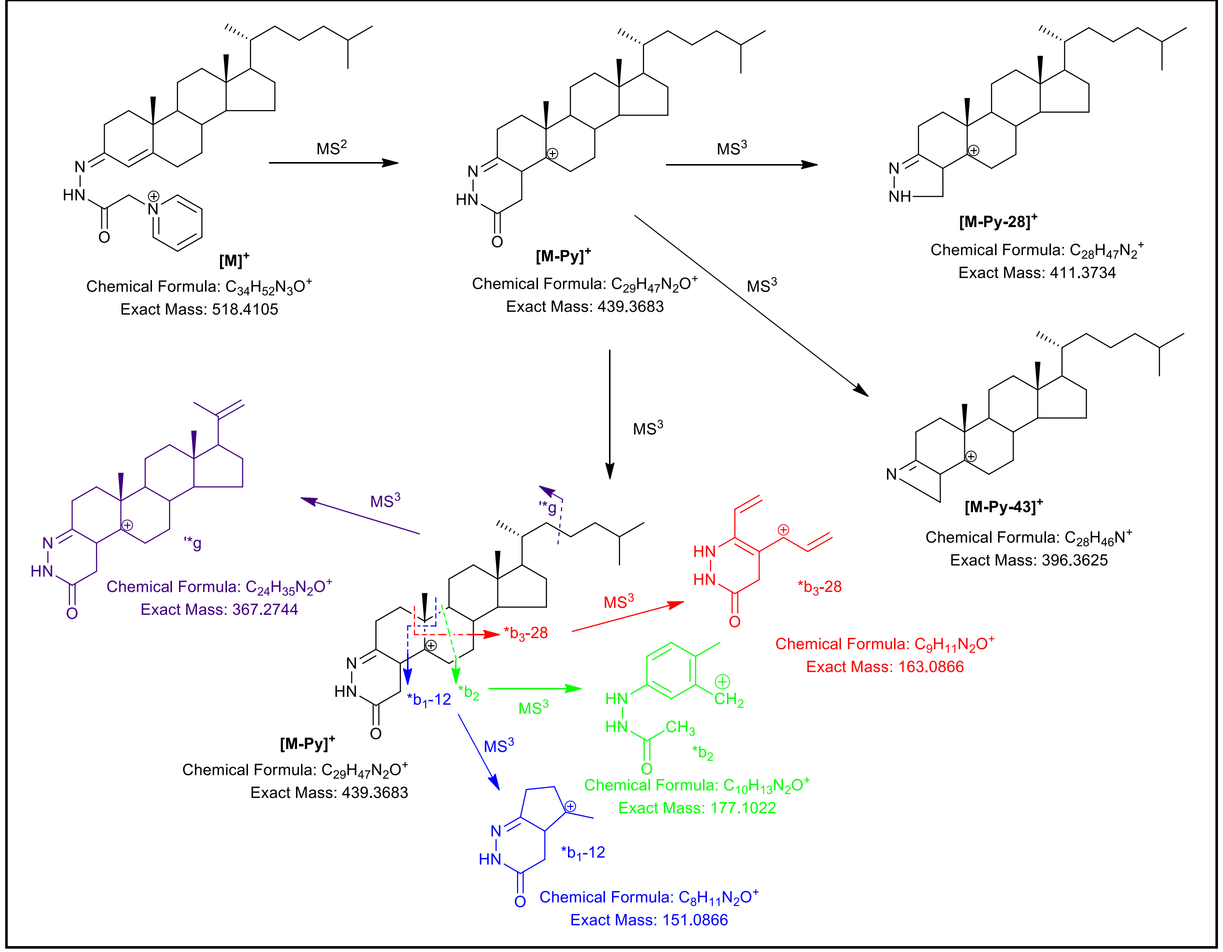

MS2
[M]+
Chemical Formula: C34H52N3O+
Exact Mass: 518.4105
[M-Py]+
Chemical Formula: C29H47N2O+
Exact Mass: 439.3683
MS3
[M-Py-28]+
Chemical Formula: C28H47N2+
Exact Mass: 411.3734
[M-Py-43]+
Chemical Formula: C28H46N+
Exact Mass: 396.3625
*g
Chemical Formula: C24H35N2O+
Exact Mass: 367.2744
*b3-28
Chemical Formula: C9H11N2O+
Exact Mass: 163.0866
*b2
Chemical Formula: C10H13N2O+
Exact Mass: 177.1022
*b1-12
Chemical Formula: C8H11N2O+
Exact Mass: 151.0866

Figure S2

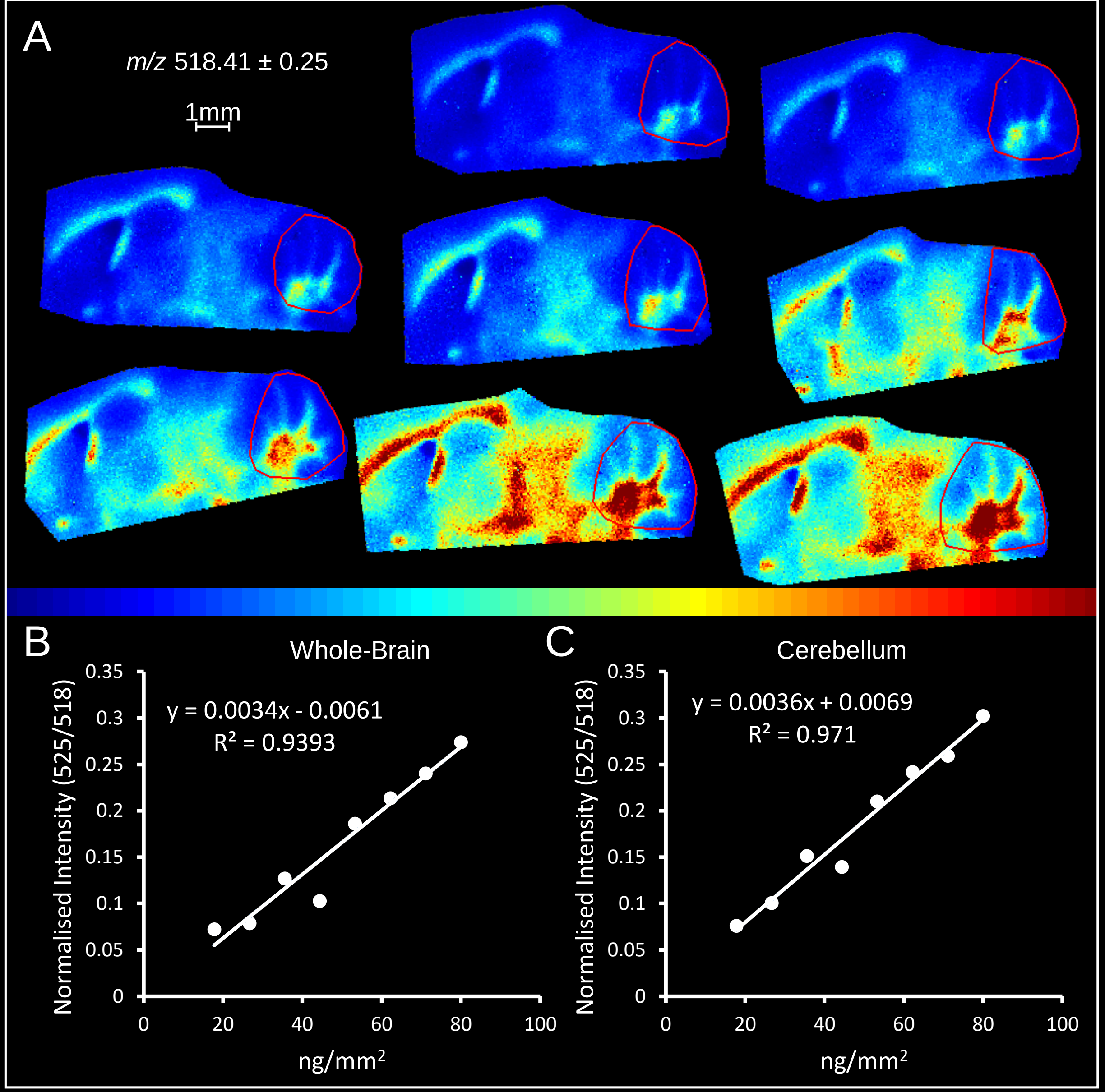
A
m/z 518.41 ± 0.25
1mm
B
Whole-Brain
y = 0.0034x - 0.0061
R² = 0.9393
Normalised Intensity (525/518)
ng/mm²
0 0.05 0.1 0.15 0.2 0.25 0.3 0.35
0 20 40 60 80 100
C
Cerebellum
y = 0.0036x + 0.0069
R² = 0.971
Normalised Intensity (525/518)
ng/mm²
0 0.05 0.1 0.15 0.2 0.25 0.3 0.35
0 20 40 60 80 100

Figure S3

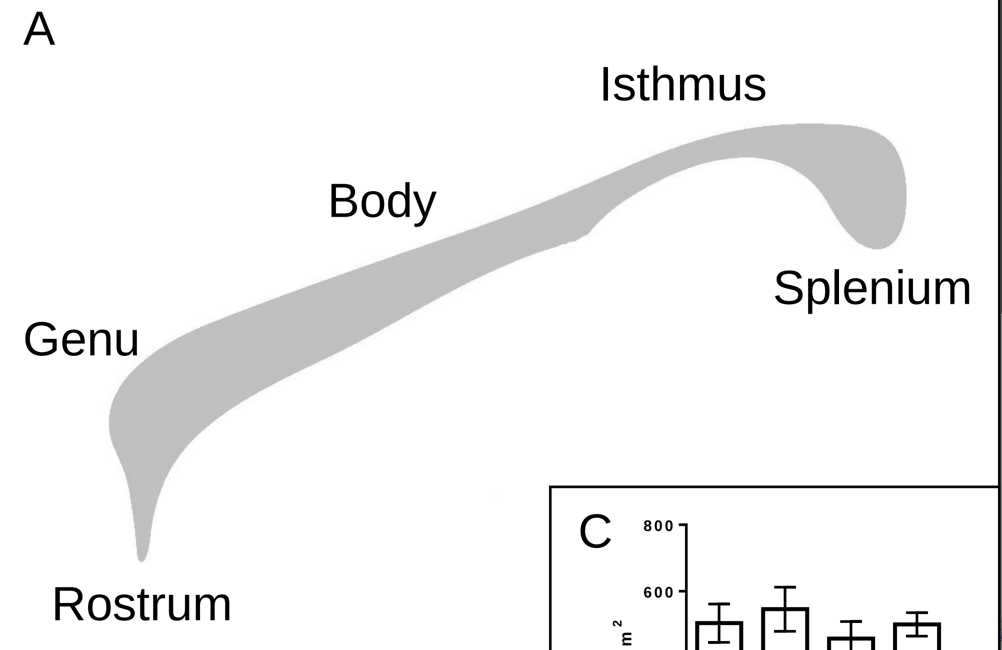


| Corpus Callosum | Cholesterol (ng/mm²) |
|---|---|
| **Rostrum-Genu** | 504.3 ± 57.7 |
| **Body** | 546.2 ± 66.3 |
| **Isthmus** | 457.8 ± 51.2 |
| **Splenium** | 500.4 ± 35.3 |

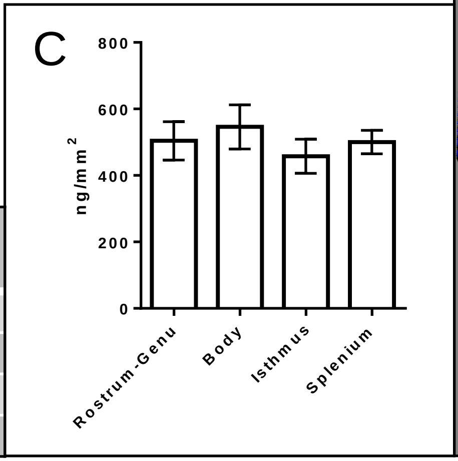


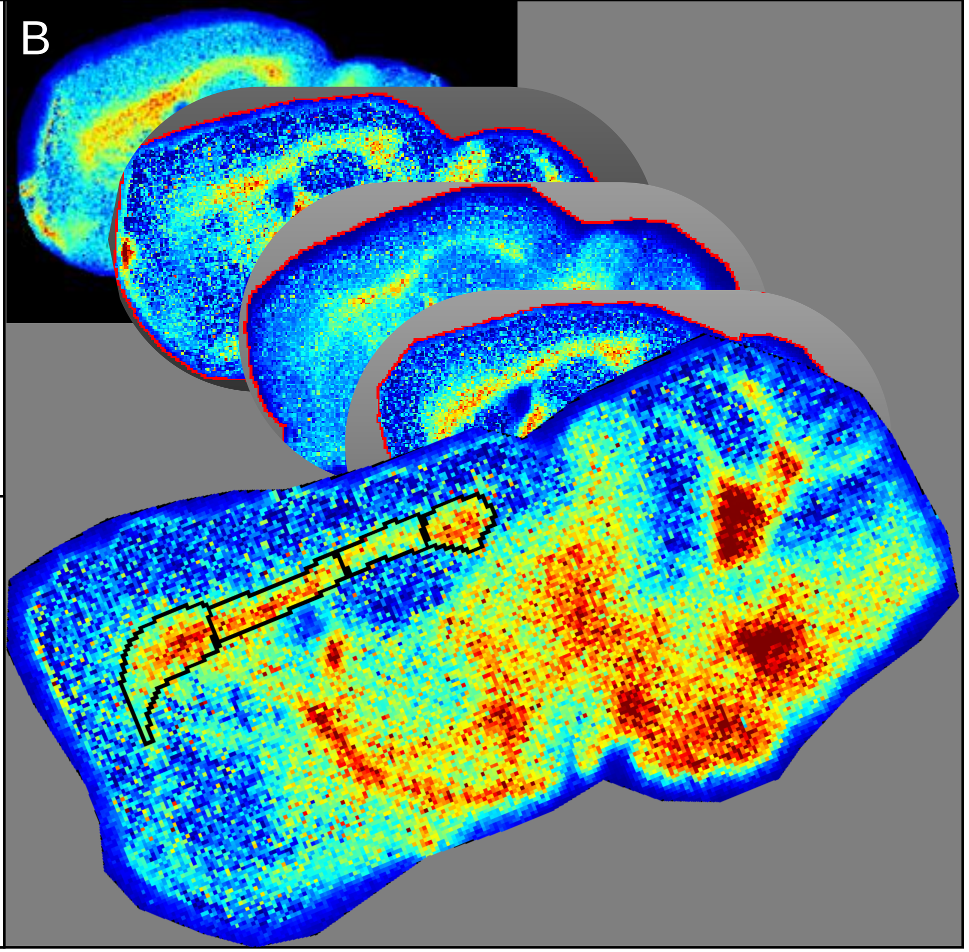

Figure S4

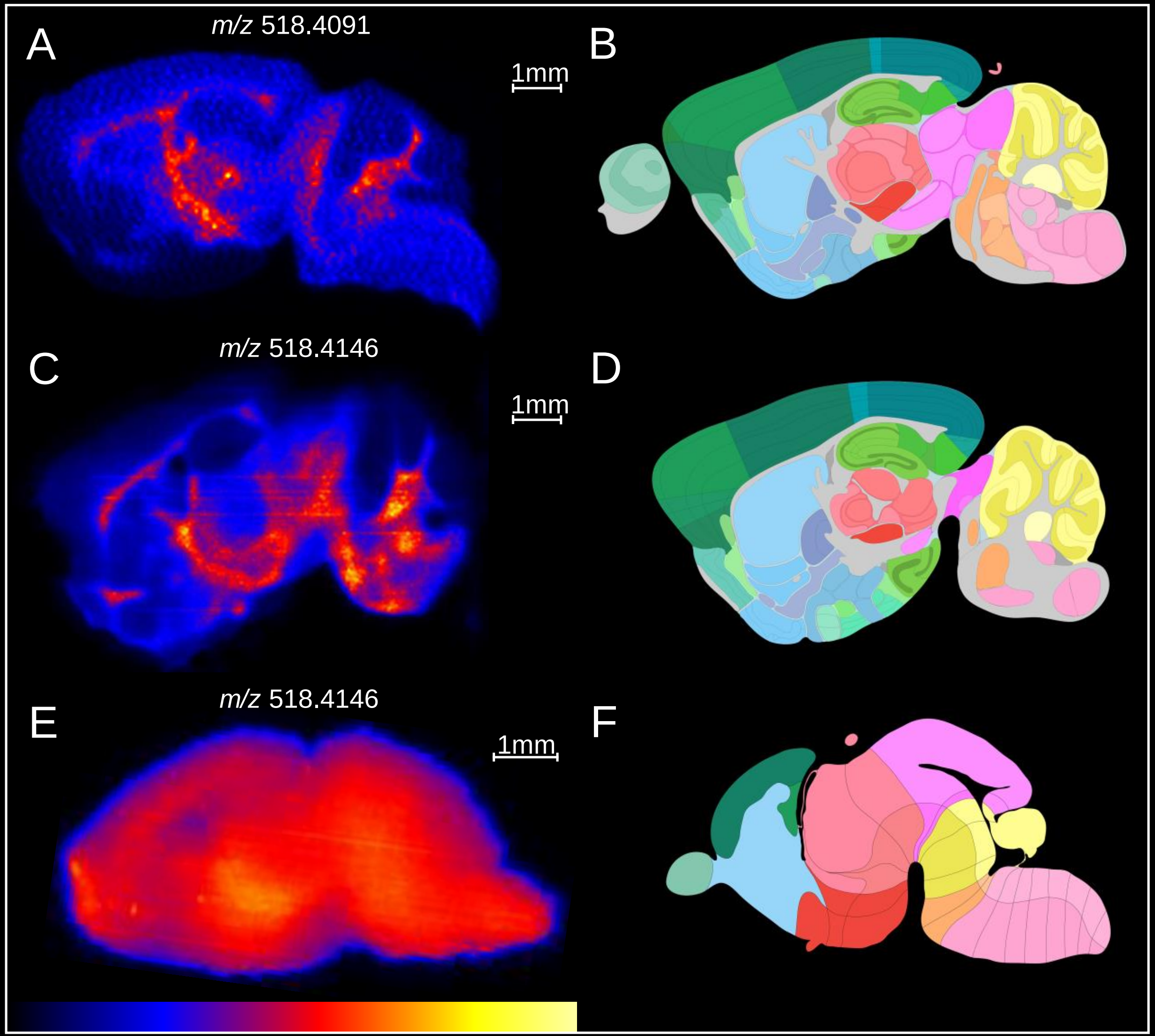

### Figure S5

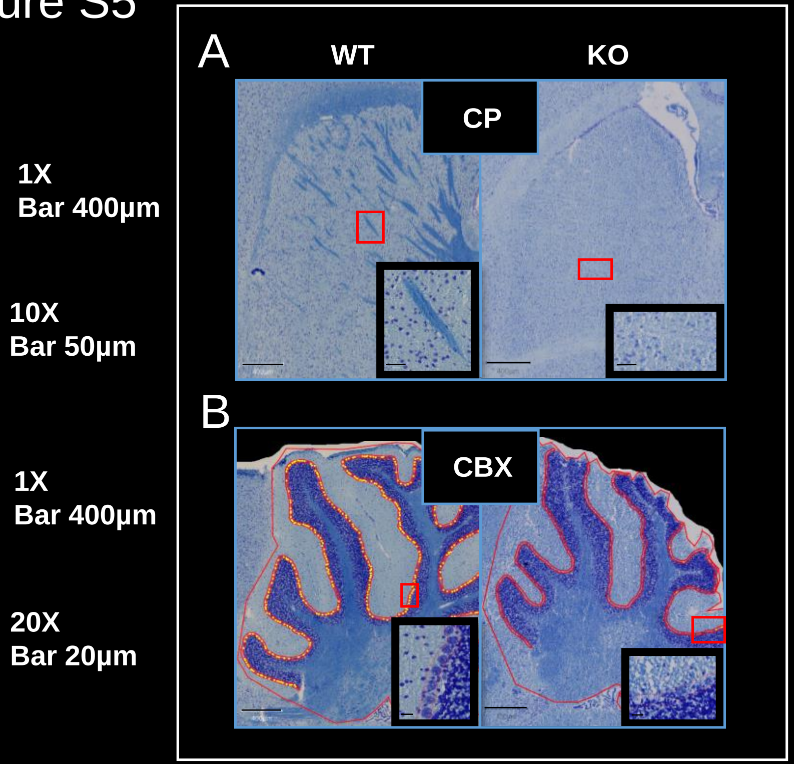
1X
Bar 400μm
10X
Bar 50μm
1X
Bar 400μm
20X
Bar 20μm
A
WT
KO
CP
B
CBX

Figure S6

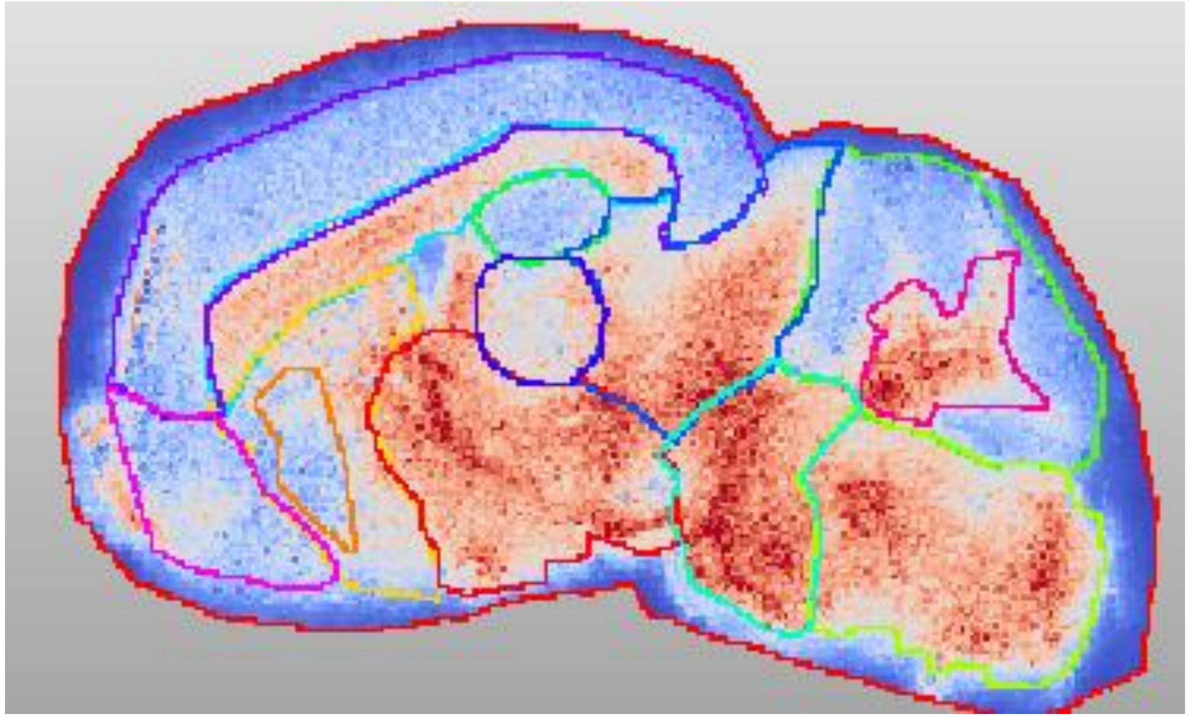
